# Site-Specific Inhibition of Translation Initiation via 2’-O-methylation

**DOI:** 10.1101/2025.06.14.659565

**Authors:** Adam Suh, Stephanie Mou, Emmely A. Patrasso, Hannah Serio, Kevin Vasquez, Rui Wang, Smriti Sangwan, Yang Yi, Daniel Arango

## Abstract

Translation initiation involves a concerted set of intermolecular interactions that efficiently recognize optimal AUG start codons. However, translation initiation complexes often start at upstream non-optimal AUGs or near-cognate codons, leading to the expression of upstream Open Reading Frames (uORFs). Using retrospective analyses of translation preinitiation cryo-EM structures, we identified putative hydrogen bonds between the 2’-OH groups of mRNA start codons with 18S rRNA. Disruption of these interactions using a chemical modification of mRNA, 2’-O-methylation (N_m_), repressed translation initiation by preventing the preinitiation complex from recognizing start codons. Notably, 2’-O-methylation in upstream AUG and near-cognate codons inhibits upstream translation initiation while enhancing the expression of canonical ORFs. These findings revealed a transcript- and site-specific inhibitory role for 2’-O-methylation in translation initiation, providing novel insights into the mechanisms of start codon selection in human transcriptomes.

## INTRODUCTION

During translation initiation, a 43S ribonucleoprotein particle containing the small ribosome subunit (40S), the initiator tRNA (tRNA_i_^met^), and translation initiation factors, binds the 5’ end of mRNA to form a 48S preinitiation complex (1). This complex scans the 5’ UTR until it encounters an optimal start codon (1). The optimality of a start codon is typically determined by the Kozak sequence (<u>R</u>CCAUG<u>G</u>, where R is either adenosine or guanosine) (2). Once the start codon is recognized, GTP hydrolysis and rearrangement of the 48S complex enable the large 60S subunit to bind and form the 80S initiation ribosome (1). Subsequently, the 80S ribosome enters elongation and continues decoding through an Open Reading Frame (ORF) until a stop codon is encountered (1).

While the Kozak sequence defines the optimal translation initiation site, translation can also initiate at suboptimal or cryptic start sites characterized by the presence of AUGs in non-Kozak sequences or near-cognate start codons, which differ from AUG by one nucleotide (CUG, GUG, UUG, AAG, ACG, AGG, AUA, AUC, and AUU) (3). While these cryptic start codons are usually bypassed by the 43S scanning complex through a process known as leaky scanning (4), translation initiation at upstream sites results in the expression of upstream Open Reading Frames (uORFs) (3). Initially thought to be mere cis-regulatory sequences that inhibit the expression of downstream canonical ORFs, recent studies demonstrated that uORFs also yield stable micropeptides with diverse biological functions, many of which are involved in various conditions such as cancer, neurodegeneration, and metabolic diseases (5–7). However, the mechanisms underlying the selection of non-canonical start codons and subsequent uORF expression in disease states remain unclear.

One poorly characterized layer of translation regulation is the epitranscriptome, defined as the set of more than 150 chemical modifications of ribonucleotides that alter RNA function. Several RNA modifications have been reported to regulate translation initiation. For instance, the hypermodified nucleotide 6-threonyl-carbamoyl-adenosine (t^6^A) in position 37 of tRNA_i_^met^ directly interacts with Kozak sequences, where it stabilizes the translation preinitiation complex (8,9). Moreover, the mRNA cap is characterized by a non-templated 7-methylguanosine (m^7^G), followed by one or two templated nucleotides with methylation in the 2’-OH of the ribose, a modification called 2’-O-methylation (also referred to as A_m_, C_m_, G_m_, U_m_, or simply N_m_) (10). When the first nucleotide is an adenosine, it is also modified into N^6^,2′-O-dimethyladenosine (m^6^A_m_) (11). The m^7^G cap is essential for translation, whereas the subsequent N_m_ modifications in the first or second nucleotide regulate mRNA stability (10–12). Beyond the cap, several mRNA modifications are found internally in the 5’UTRs, including m^6^A (13–15), N4-acetylcytidine (ac^4^C) (8,16), and 2′O-methylation (17–20). While m^6^A regulates cap-independent translation by recruiting ‘reader’ proteins (10–12), and ac^4^C weakens the interaction of tRNA_i_^met^ with the Kozak sequence (8), the function of internal 2′O-methylation sites within 5’UTRs is unknown.

Using synthetic mRNAs with 2′-O-methylation at every position of the Kozak sequence, we found that N_m_ at the first nucleotide of the AUG, CUG, ACG, and AUC start codons inhibits translation initiation. Mechanistically, 2’-O-methylation prevents the preinitiation complex from recognizing start codons. Notably, N_m_ at upstream start codons inhibits upstream initiation while enhancing translation from downstream canonical AUGs. Thus, this study revealed an inhibitory effect of 2’-O-methylation in start codon recognition, providing novel insights into the mechanisms of start codon selection in human cells. This study also provides the molecular basis for the potential use of 2’-O-methylation to inhibit aberrant uORF expression or enhance downstream ORF expression during RNA therapeutics.

## MATERIALS AND METHODS

### Cell culture

HEK293T cells were cultured in Dulbecco’s Modified Eagle Medium (DMEM, Gibco, Cat. #: 11965092) supplemented with 10% Fetal bovine serum (Corning, Cat. #: 30-015-CV) and 1% Penicillin/Streptomycin (Dot scientific, Cat. #: DSP93560-50). HeLa cells were cultured in DMEM (Gibco, Cat. #: 11965092) supplemented with 10% bovine calf serum (Cytiva, Cat. #: SH30073.04) and 1% Penicillin/Streptomycin. Cells were grown at 37 °C in a humidified environment with 5% CO_2_.

### Synthesis and capping of 5’ RNA fragments

To produce the 170 nt 5’UTR, an antisense DNA template containing a T7 promoter sequence was purchased from Integrated DNA Technologies (IDT) and annealed to a sense T7 primer (Supplementary Table S1). To produce the 504 nt 5’UTR, we generated the DNA template by PCR using primers that annealed to a plasmid containing the coding sequence of the micro red fluorescence protein, ensuring that the amplicon excluded any start codon. The forward primer contained the T7 sequence as an overhang. PCR was performed using the Q5 Hot Start High-Fidelity 2X Master Mix (New England Biolabs, Cat. #: M0494).

The 170 nt and 504 nt 5’UTRs were *in vitro* transcribed using the HiScribe® T7 High Yield RNA Synthesis Kit (New England Biolabs, Cat. #: E2040), following the manufacturer’s suggestions. RNA was purified using 1.8X SPRI beads (CleanNGS Cat. #: CNGS-0001). Purified RNA was capped *in vitro* using the Vaccinia Capping System (New England Biolabs, Cat. #: M2080) and re-purified using 1.8X SPRI beads. The capped 27 nt 5’UTR was custom-synthesized by TriLink BioTechnologies.

The integrity of RNA fragments was verified using polyacrylamide gels prepared in a tris-borate-urea buffer (2 mL of 40% PAA [19:1], 5 g of Urea, 1mL of 10X TBE, complete to 10 mL ddH2O). Electrophoresis was run in a 0.5X TBE running buffer and stained with a 1X SYBR gold solution (ThermoFisher, Cat. #: S11494). Gels were visualized in a Sapphire Biomolecular Imager (Azure Biosystems). Primers, RNA oligonucleotides, and resulting DNA sequences are described in Supplementary Table S1.

### Dot blots

Dot blots were performed to verify the capping of *in vitro* transcribed RNAs. Briefly, 500 ng RNA was denatured at 65°C for 5 min and loaded onto Hybond-N+ membranes. Membranes were crosslinked with 120 mJ^2^/cm^2^ in the UV Crosslinker. Membranes were first stained in a 0.2% methylene blue solution (0.2% methylene blue in 0.4M sodium acetate and 0.4M acetic acid), then washed three times in DI water, and imaged in a Sapphire Biomolecular Imager.

Membranes were then blocked with 1% bovine serum albumin (BSA) in PBS containing 0.05% Tween-20 (PBST) for 15 min at room temperature and probed overnight with mouse anti-m^7^G antibodies (1:1000, MBL International Corporation, Cat. #: RN016M) in 1% BSA at 4°C. Membranes were next washed three times with 0.1% PBST, incubated with HRP-conjugated secondary anti-mouse IgG (1:10000 dilution, Cell Signaling Technologies, Cat. # 7076S) in 1% BSA at room temperature for 1 hr, washed three times with 0.1% PBST and developed with enhanced chemiluminescent (ECL) western blotting substrate (Promega, Cat #: W1001) on a Sapphire Biomolecular Imager.

### Synthesis and polyadenylation of 3’ RNA fragments

To produce the ORF and 3’UTR of NanoLuc, we generated a DNA template by PCR using primers that annealed to a plasmid containing the NanoLuc coding sequence, ensuring the amplicon excluded the canonical start codon. The forward primer contained the T7 sequence as an overhang. To produce the ORF and 3’UTR of GFP, we generated a DNA template by PCR using primers that annealed to a plasmid containing the GFP coding sequence, ensuring the amplicon excluded the canonical start codon. The forward primer contained the T7 sequence as an overhang. PCR was performed using the Q5 Hot Start High-Fidelity 2X Master Mix (New England Biolabs, Cat. #: M0494). To produce the mRNA reporter containing a uORF, the canonical NanoLuc ORF, and the 3’UTR, we generated the DNA template by PCR using primers that annealed to a plasmid that incorporated an uORF or an overlapping uORF upstream of the coding sequence of NanoLuc, making sure to exclude the upstream canonical start codon from the amplicon product. Primers and resulting DNA sequences are described in Supplementary Table S1.

DNA duplexes were *in vitro* transcribed using the HiScribe® T7 High Yield RNA Synthesis Kit (New England Biolabs, Cat. #: E2040) according to the manufacturer’s instructions. RNA was purified using 1X SPRI beads (CleanNGS Cat. #: CNGS-0001). RNA fragments were dephosphorylated with 100 U of calf intestinal alkaline phosphatase (CIAP, Promega, Cat. #: M2825), followed by mono-phosphorylation of the 5’ end using 15 U of T4 Polynucleotide Kinase (PNK, New England Biolabs, Cat. #: M0201L) in the presence of ATP. Alternatively, *in vitro* transcribed RNAs were treated with RppH (New England Biolabs, Cat. #: M0356S), which converts 5’-triphosphates into 5’-monophosphates. 5’-monophosphorylated RNA products were purified using the RNA Clean & Concentrator-25 Kit (Zymo Research, Cat. #: R1018). RNA products (10 μg) were polyadenylated using 5 U of *E. coli* Poly(A) Polymerase (New England Biolabs, Cat. #: M0276). The integrity of polyadenylated RNA fragments was verified using TBU-PAGE. Electrophoresis was run in 0.5X TBE buffer and stained with 1X SYBR Gold.

### Splint ligations

5’-monophosphorylated RNA oligonucleotides (12 nt) containing the Kozak sequence with an AUG, CUG, GUG, ACG, or AUC start codons were purchased from IDT. Likewise, the 2’-O-methylated versions were purchased from IDT (Supplementary Table S1). For three-way splint ligations, these RNA oligos were combined with 5’ and 3’ *in vitro* transcribed RNA fragments at equimolar concentrations (15 pmol). An antisense DNA oligonucleotide (15 pmol) was added to splint the different RNA pieces. Reactions were denatured at 65 °C for 2 minutes and ramped down to room temperature for annealing. For two-way splint ligation, unmethylated or 2’-O-methylated capped RNA oligonucleotides were purchased from TriLink BioTechnologies and combined with a 3’ *in vitro* transcribed RNA fragment at equimolar concentrations (15 pmol). An antisense DNA oligonucleotide (15 pmol) was added to splint the different RNA pieces.

Ligations were performed using 10 U of T4 RNA ligase 2 (dsRNA ligase, New England Biolabs, Cat. #: M0239) for 4 hr at 37 °C, according to the manufacturer’s recommendations. Reactions were stopped with 2 U of Turbo DNase I (ThermoFisher Scientific, Cat. #: AM2238) for 30 min at 37°C and purified using the RNA Clean & Concentrator-5 kit (Zymo Research, Cat. #: R1014). To verify the integrity of splint-ligated NanoLuc RNA, 500 ng of ligated RNAs were separated by 4% TBU-PAGE and stained with 1X SYBR gold (Invitrogen, Cat. #: S11494). To verify the integrity of splint-ligated GFP RNA, 500 ng of ligated RNAs were separated by 1% denaturing agarose gels and stained with 1X SYBR gold (Invitrogen, Cat. #: S11494). Gels were visualized in a Sapphire Biomolecular Imager (Azure Biosystems).

To purify ligated RNA fragments, gels were stained with SYBR Gold, imaged, and aligned to a printed scan to guide precise excision of RNA bands. Target gel slices were excised, crushed, and RNA was eluted overnight at room temperature in elution buffer (300 mM NaCl, 10 mM Tris pH 7.0, 1 mM EDTA, 0.25% SDS). Eluates were clarified by centrifugation, and RNA was precipitated with glycogen and ethanol at −80 °C, followed by centrifugation. Pellets were washed twice with 70% ethanol, air-dried, and resuspended in nuclease-free water. The integrity of purified mRNA reporters was verified by TBU-PAGE and SYBR gold staining.

### NanoLuc and Firefly luciferase assays

HEK293T or HeLa cells were seeded at a density of 30,000 cells in 100 μL complete DMEM medium in 96-well plates. Cells were cultured for 24 hr before transfection with 300 ng of capped and polyadenylated splint-ligated RNA or an unligated RNA control. Transfection reactions were prepared using the TransIT-mRNA Transfection Kit (Mirus, Cat. #: MIR2225) following the manufacturer’s instructions. Following the addition of the transfection mixtures, cells were incubated for 6 hrs at 37°C. For luminescence detection, cells were assayed with Nano-Glo® Luciferase Assay System (Promega, Cat. #: N1110) according to the manufacturer’s instructions for 96-well plates. For the dual transfection experiments, cells were assayed using the Nano-Glo Dual-Luciferase Reporter Assay System (Promega, Cat. #: N1610) according to the manufacturer’s instructions for 96-well plates. Luminescence was read in a BioTek Synergy LX Multimode Reader (Agilent Technologies).

### Flow cytometry

HEK293T or HeLa cells were seeded at a density of 150,000 cells in 1 mL medium in 12-well plates. Cells were transfected with 600 ng of capped and polyadenylated splint-ligated mRNA or unligated RNA control. Transfection reactions were prepared using the TransIT-mRNA Transfection Kit (Mirus) following the manufacturer’s instructions for 12-well plates. Following the addition of the transfection mixtures, cells were incubated for 18 hrs at 37°C. Cells were trypsinized and washed with PBS three times before they were resuspended in 500 μL PBS and immediately analyzed via Flow cytometry in an LSR Fortessa 1 Analyzer or a BD FACSymphony A5-Laser Analyzer. The percentage of GFP-positive cells was determined using FlowJo v10.10.0.

### RNA isolation and RT-qPCR

RT-qPCR was performed to evaluate the efficiency of ligation before transfection or the stability of ligated mRNA at different time points (0, 3, and 6 hr) post-transfection. Briefly, total RNA was isolated from cells at 0, 3, and 6 hr post-transfection using the Trizol method. The extracted RNA was reverse transcribed using SuperScript™ IV Reverse Transcriptase (ThermoFisher Scientific, Cat. #: 18090010) using gene-specific RT primers for NanoLuc, GFP, and 18S rRNA (Supplementary Table S1). qPCR was performed using the 2X PowerUp SYBR Master Mix (ThermoFisher Scientific, Cat. #: A25742) and primers that amplify the ligated mRNAs, total mRNAs (ligated plus unligated), and 18S rRNA (Supplementary Table S1). To obtain the efficiency of ligation, we applied the formula 100 * 2^[-(Ct^ligated^/Ct^total.mRNA^). To measure mRNA stability, we applied the formula final.time(2^[-(Ct^ligated^/Ct^18S.rRNA^)/initial.time(2^[-(Ct^ligated^/Ct^18S.rRNA^).

### *S*ucrose density gradients

Cy5-labeled synthetic mRNAs were purchased from Genescript (Supplementary Table S1). Reticulocyte lysates (37.5 μL) were pre-incubated with vehicle DMSO, 1 mM GMPPNP (Guanosine 5′-[β,γ-imido]triphosphate trisodium salt hydrate, Millipore Sigma, Cat. #: G0635), or 100 μg/ml Harringtonine (Abcam, Cat. #: ab141941) prior to adding 150 nM of Cy5-labeled synthetic mRNAs in a total volume of 50 μL. Reactions were incubated for 10 min at 30 °C and immediately stopped on ice. Reactions were loaded on a 5-30% sucrose density gradient (50 mM Tris-acetate [pH 7.0], 50 mM NH_4_Cl, 12 mM MgCl_2_, 1 mM DTT) poured using a BioComp Gradient Master. Gradients were spun at 42,000 rpm in a Beckmann Coulter SW41 Ti rotor for 2:26 hr at 4°C. Recording of Cy5 fluorescence in different fractions was performed using a BioComp Density Gradient Fractionation system.

### siRNA transfection

The siRNAs used in this study were purchased from Thermo Fisher (siFBL-1 Cat. #: s4820; siFBL-2 Cat. #: s4821; siCtrl Cat. #: 4390844) (Supplementary Table S1). The siRNAs were transfected into HeLa cells using the Lipofectamine RNAiMAX Transfection Reagent (ThermoFisher Scientific, Cat. #: 13778150) by following the manufacturer’s instructions. Total RNA and cell lysates were collected three days post-transfection. The efficiency of FBL knockdown was verified by Western blot.

### Western blot

Cells were washed with cold 1x phosphate-buffered saline (PBS, Corning, 21-040-CM) and lysed in RIPA lysis buffer containing 150mM NaCl, 1% Triton X-100, 0.5% sodium deoxycholate, 0.1% SDS, and 50mM Tris (pH 8.0). Lysates were centrifuged at 13,200 rpm for 10 minutes at 4 °C, and protein concentrations were determined using Micro BCA^TM^ Protein Assay Kit (ThermoFisher Scientific, 23235). Thirty micrograms of protein were separated by 12% SDS-PAGE Tris-Glycine at 100 V for 1.5 hours and transferred overnight at 4 °C to a nitrocellulose membrane. Membranes were blocked with 5% non-fat milk in Tris-buffered saline with 0.05% Tween-20 (TBST) for at least 15 minutes at room temperature. Membranes were then incubated overnight at 4 °C with one of the following primary antibodies in 1% non-fat milk in TBST: rabbit polyclonal anti-FBL (1 microgram/mL, Cat. #: ab5821, abcam); rabbit polyclonal anti-β-actin (1:2000, Cat. #: ab8227, abcam); mouse monoclonal anti-Nucleostemin (anti-GNL3) (1:1000, Cat. #: sc-166460, Santa Cruz Biotechnology); mouse monoclonal anti-Nek2 (1:1000, Cat. #: sc-55601, Santa Cruz Biotechnology); or mouse monoclonal anti-SF2/ASF (anti-SRSF1) (1:1000, Cat. #: sc-33652, Santa Cruz Biotechnology). Membranes were washed three times (if monoclonal) or five times (if polyclonal) with TBST and incubated with HRP-conjugated anti-rabbit IgG (1:5000, Cat. #: 7074, Cell Signaling Technology) or HRP-conjugated anti-mouse IgG (1:5000, Cat. #: 7076, Cell Signaling Technology) for 1 hour at room temperature. Signals were detected using ECL Western Blotting Substrate (Promega, W1001) or SuperSignal^TM^ West Femto Maximum Sensitivity Substrate (ThermoFisher Scientific, 34095) and visualized with a Sapphire multimode imager (Azure Biosystems). Band densitometry was quantified using ImageJ. The relative expression levels were estimated by normalizing to the β-actin loading control.

### Identification of 2’-OH hydrogen bonds in translation preinitiation complexes

The following structures were downloaded from the Protein Data Bank: 7UCJ (rabbit reticulocyte), 6YAL (rabbit reticulocyte), 8PJ1-6 (reconstituted human 48S). Structural visualization was performed in PyMOL. Briefly, the background was set to white, and all elements were initially hidden. Specific residues of the mRNA, centered around nucleotides N[-3], N[-2], N[-1], N[+1], N[+2], N[+3], and N[+3] were selected and visualized as both ribbons (with increased width) and sticks. Specific residues within tRNA_i_^met^, 18S rRNA, eIF1A, and eIF2α that interact with mRNA were also selected. Hydrogen bond calculations were performed between mRNA, tRNA_i_^met^, 18S rRNA, eIF1A, and eIF2α. Hydrogen-bond geometry involving the ribose O2′ group, distances between O2′–P and O2′–O2 atoms, and corresponding heavy-atom proxy angles were measured using PyMOL. Density maps and corresponding models were visualized in ChimeraX. Molecular animation videos were generated using the movie-recording tools. Contour levels were set below the EMDB-recommended values for all except 7UCJ and 6YAL, for which the levels were adjusted to show the density clearly. Supplementary Video 1_7UCJ contour levels = 0.0183 emdb_26444, Supplementary Video 2_6YAL contour levels = 0.0196 emdb_10760, Supplementary Video 3_7UCJ contour levels = 0.016 emdb_26444, Supplementary Video 4_8PJ3 contour levels = 5.82 emdb_17698, Supplementary Video 5_8PJ3 contour levels = 4.0 emdb_17698, Supplementary Video 5_8PJ3 contour levels = 4.73 emdb_17698.

### Identification of N_m_ sites in AUG and near-cognate codons

Maps of 2’-O-methylation in C4-2 and HeLa cells were extracted from Li et al. 2024 (17). N_m_ sites with a 2’-O-methylation ratio above ten percent were selected for downstream analysis. To identify N_m_ sites in canonical AUG codons, we extracted the genomic information for Ensembl canonical start codons from a human genome annotation file (GRCh38). Then, bedtools/2.29.2 was used to identify N_m_ sites around canonical start codons. N_m_ sites were then binned by their location relative to the canonical AUG, selecting only sites at position N_m_[+1]. To identify N_m_ sites in upstream AUG and near-cognate codons, we selected N_m_ sites within protein-coding genes that aligned to 5’UTR in any possible transcript. Then, we extracted the genomic sequence flanking each N_m_ site. We filtered for sequences containing AUG and near-cognate codons. Each N_m_ site location was derived relative to the AUG or near-cognate codons N_m_[+1]. This analysis was performed using custom scripts in Biopython 1.85.

### RTL-P assay

To confirm the N_m_ occurrence along with its alteration upon FBL depletion, the RTL-P assay was conducted as previously described (17). First, total RNAs from control and FBL-knockdown HeLa cells were isolated using the RNeasy Plus Mini Kit (Qiagen). Reverse transcription (RT) was then performed using specific reverse primers targeting mRNA sequences downstream of the 2’-O-methylation site in the presence of either a low (1 μM) or a high (1 mM) concentration of dNTPs. The obtained cDNA was subjected to two separate amplification reactions using one pair of primers targeting upstream and downstream of the methylation site, and another pair targeting a downstream region that would not be interrupted by the 2’-O-methylation site (Supplementary Table S1). The PCR mixture consisted of 10 μL AmfiSure PCR master mix (GenDEPOT), 2 μL cDNA, and 1 mM PCR primers. The PCR products were then separated using 1% agarose gels and photographed using a Bio-Rad imaging system. DNA band signal intensities were calculated using ImageJ. The methylation ratio of each group was determined from the densities of the PCR bands obtained using high and low dNTP concentrations. The RT and PCR primers designed for the RTL-P assay are listed in Supplementary Table S1.

### Codon bias analysis

Using a list of genes that were expressed in HeLa HR-Ribo-seq (8), 5’UTR sequences of the longest transcript per gene were extracted from Ensembl/113 with BioMart. This was also performed for genes containing N_m_[+1]. The frequencies of AUG and near-cognate codons were calculated for modified codons in the 5’UTRs and for the presence of these codons in the 5’UTRs of expressed protein-coding genes. Codon frequencies were analyzed using a chi-square test of proportions with a Benjamini-Hochberg correction.

### Gene ontology analysis

Genes containing N_m_[+1] in upstream AUG or near-cognate codons in their 5’UTR were used for gene ontology (GO) analysis for three ontologies: biological process, cellular component, and molecular function. An R script was used to perform and visualize this analysis using the clusterProfiler/4.14.6 and org.Hs.eg.db/3.20.0 packages. The top 5 categories for each ontology were displayed in descending order of *p-value*.

### Analysis of translation efficiency in HeLa cells

We analyzed publicly available Ribo-seq performed in HeLa cells (GSE105248) (21). This dataset contained three Ribo-seq and Input RNA-seq from FBL knockdown (Doxycycline-inducible shRNA against FBL) and control groups. Ribosome-protected fragments (RPFs) were derived by trimming adapters from raw sequence files with TrimGalore/0.6.10, aligning to the transcriptome with STAR/2.7.9a, and featureCounts extracted RPFs with subread/2.0.3. mRNA and RPF counts were normalized with DESeq2/1.46.0, and translation efficiency was calculated as normalized RPF counts divided by normalized mRNA counts. We included only protein-coding genes with normalized mRNA counts> 0. The log2FoldChange of translation efficiency was calculated with Rstudio/v4.4.0. We compared genes with mapped N_m_[+1] at an upstream AUG or near-cognate codon in the 5’UTR with those with no mapped N_m_ in the 5’UTR in HeLa cells. Differences in translation efficiency between groups were analyzed with a Wilcoxon rank-sum test.

### Codon usage in upstream initiation sites

Codon identity of translation initiation sites from Harringtonine Ribo-seq in HeLa cells was downloaded from Arango et al., 2022 (8). Data were binned by codon identity and visualized as a pie chart in RStudio/v4.4.0 using the packages ggpubr and ggplot2.

### Other statistical analysis

All other statistical tests and plotting were performed in Rstudio/v4.4.0. The specific statistical tests, graph type, variables, number of observations per variable, and *p-values* are indicated in the figures or their legends.

## RESULTS

### 2’-OH hydrogen bonds within translation initiation complexes

To understand what defines an optimal start codon, several groups solved cryo-EM structures of mammalian preinitiation and initiation complexes (8,9,22,23). These efforts provided a snapshot of the crucial intermolecular interactions surrounding Kozak sequences (Fig. 1A). Besides the recognition of start codons by the anticodon loop in tRNA_i_^met^, the interactions of positions A/G[-3] and G[+4] with translation initiation factors are required for translation initiation (Fig. 1A). In revisiting several mammalian translation initiation structures (8,9,22), we observed that 2’-OH groups of mRNA nucleotides within the Kozak sequence appeared to be interacting with other components of the preinitiation complex. For instance, t^6^A in position 37 of tRNA_i_^met^ is in proximity with the 2’-OH group of the nucleotide preceding the start codon, C[-1] (Fig. 1B, Supplementary Fig. S1A, and Supplementary Table S2). The 2’-OH group of the first nucleotide, A[+1], in the start codon appears to interact with U1830 of 18S rRNA, with the phosphate (PO_4_^3-^) in the backbone (Fig. 1C), and possibly with the keto (C=O) group in position C2, especially in the late stages of the preinitiation complex (Fig. 1D and Supplementary Fig. S1B-F). The H-bonds between the 2’-OH groups in C[-1] and A[+1] with tRNA and 18S rRNA, respectively, are supported by structural models, electron densities, distances between O2′–P and O2′–O2 atoms, and the corresponding heavy-atom proxy angles (Fig. 1B and C, Supplementary Videos 1-4, and Supplementary Table S2).

**Figure 1.**
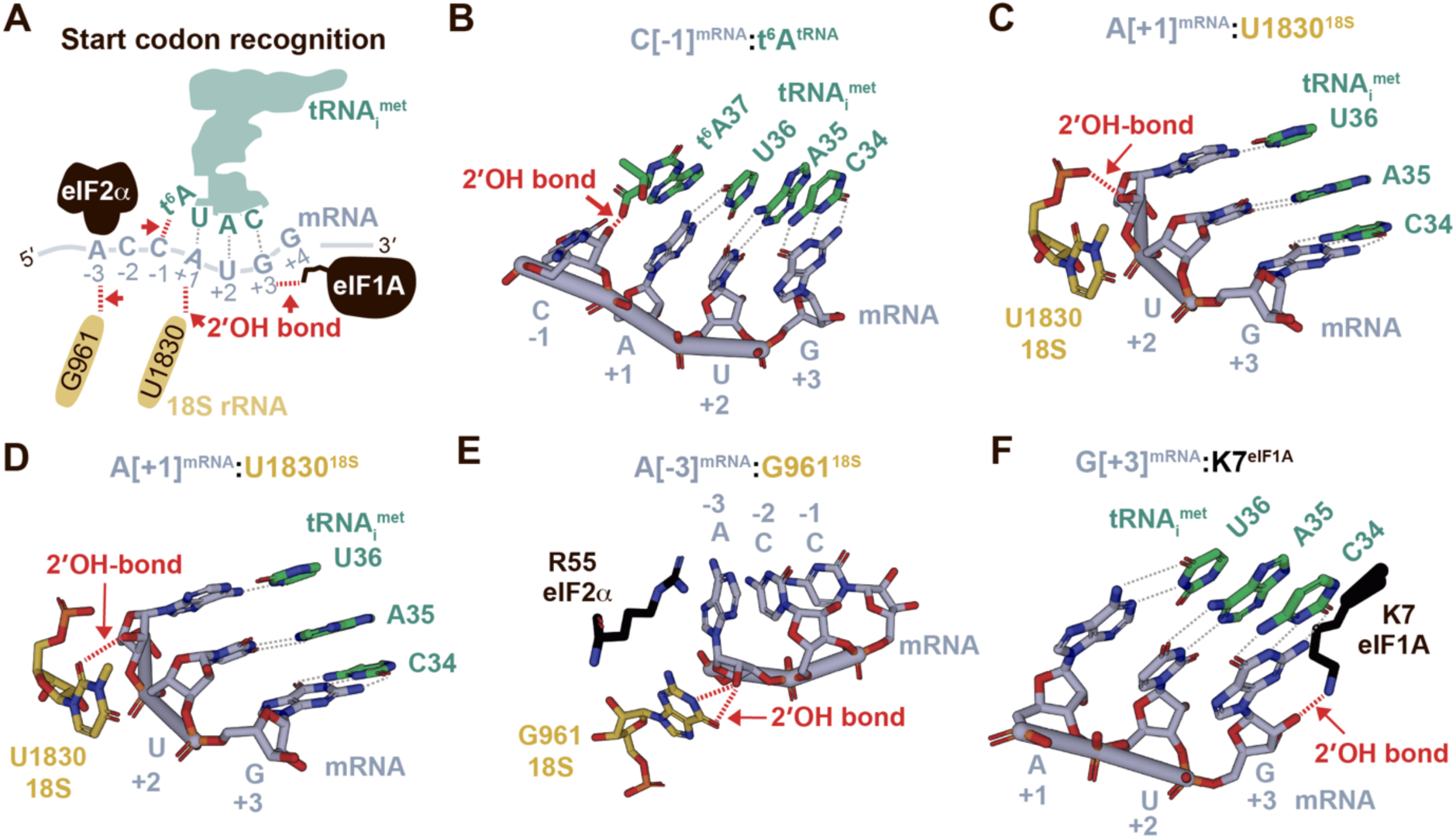
2’-OH hydrogen bonds within translation initiation complexes. **(A)** schematic representation of the intermolecular interactions within translation preinitiation complexes. (**B-D)** Cryo-EM visualization of rabbit 80S initiation ribosomes captured in complexed with harringtonine (PDB:7UCJ, Arango et al., 2022) (8) and centered around the interaction of the 2’-OH group in nucleotide C[-1]^mRNA^ with t^6^A in tRNA_i_^met^ (B); centered around the interaction of the 2’-OH group in nucleotide A[+1]^mRNA^ with the PO_4_^3-^ group of U1830 in 18S rRNA (C); or centered around the interaction of the 2’-OH group in nucleotide A[+1]^mRNA^ with the C[2]=O group of U1830 in 18S rRNA (D). (**E-F)** cryo-EM visualization of human 48S preinitiation complexes (PDB:8PJ3 Petrychenko et al., 2025) (22) centered around the interaction of the 2’-OH group in nucleotide A[-3]^mRNA^ with G961 in 18S rRNA (E); or centered around the interaction of the 2’-OH group in nucleotide G[+3]^mRNA^ with lysin 7 in eIF1A (F). All structures were downloaded from PDB and visualized using PyMOL.

Additional putative H-bonds were observed between the 2’-OH group of nucleotide A[-3] in the mRNA with G961 of 18S rRNA and between the 2’-OH group of G[+3] with lysine seven (K7) of the translation initiation factor 1A (eIF1A), although the latter are not well supported by electron densities or heavy-atom proxy angles (Fig. 1E-F, Supplementary Videos 5-6, and Supplementary Table S2). Together, these observations suggest that 2’-OH groups in mRNA interact with 18S rRNA and tRNA_i_^met^, potentially playing a crucial role in start codon recognition.

### 2’-O-methylation in position A[+1] of AUG start codons inhibits translation

To probe the significance of the putative hydrogen bonds formed by 2’-hydroxyl groups in mRNA, we sought to systematically introduce a 2’-O-methylation modification in every nucleotide within the Kozak sequence, as the 2’-OCH_3_ group lacks the donor capacity of 2’-OH (Fig. 2A). While 2’-O-methylation can be incorporated at specific positions within RNA molecules by chemical synthesis, this approach is limited by the length of RNA fragments that can be chemically synthesized. We therefore implemented a three-way splint ligation approach to introduce 2’-O-methylation (Fig. 2B) (24). Briefly, a 5’UTR lacking a start codon was transcribed and capped *in vitro* (Supplementary Fig. S2A and B). In parallel, the ORF of NanoLuciferase (NanoLuc)—also lacking a start codon—was transcribed and polyadenylated *in vitro* (Supplementary Fig. S2C). These pieces were ligated to a chemically synthesized RNA fragment containing the unmodified or 2’-O-methylated Kozak sequence using a DNA splint that hybridizes to the three independent RNA pieces (Fig. 2B).

**Figure 2.**
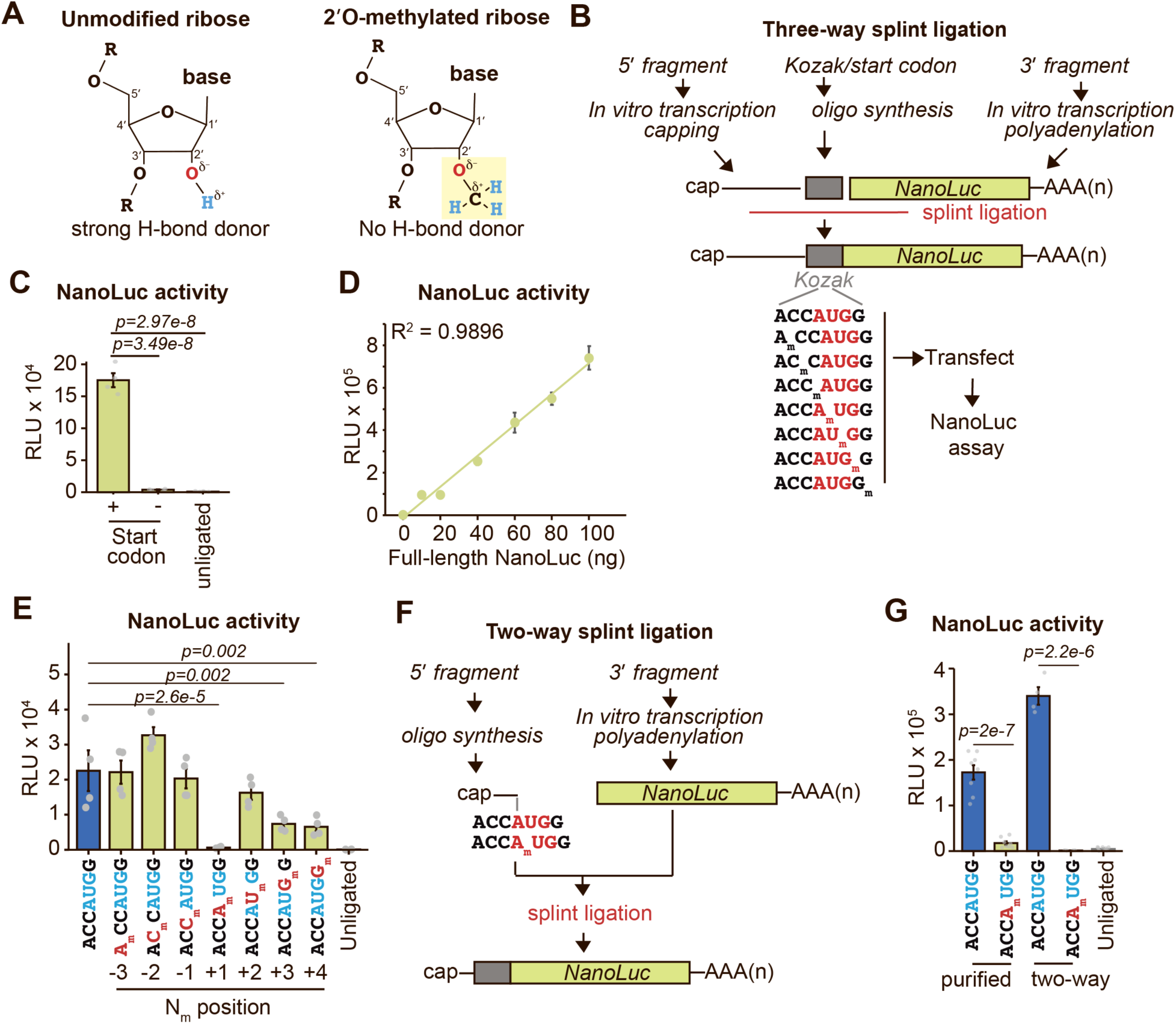
2’-O-methylation of A[+1] within AUG start codons inhibits protein synthesis. **(A)** chemical structures of unmethylated and 2’-O-methylated nucleosides. (**B)** experimental design of a three-way splint ligation approach to insert 2’-O-methylation in single positions within mRNA start codons. **(C)** relative light units (RLUs) were measured in cell lysates from HEK293T cells transfected with ligated NanoLuc mRNAs, with or without a start codon, or transfected with unligated mRNAs as a control. Mean ± SEM, n = 3. p = One-way ANOVA with Tukey’s post hoc analysis. **(D)** RLUs were measured in lysates from HEK293T cells co-transfected with increasing amounts of full-length NanoLuc mRNA reporters and constant amounts (50 ng) of unligated mRNA. Mean ± SEM, n = 3. A linear regression demonstrates the linearity of the assay. **(E)** RLUs were measured in lysates from HEK293T cells transfected with ligated NanoLuc mRNAs carrying 2’-O-methylation in different positions around the start codon. Unmethylated (blue) and unligated mRNAs were used as controls. Mean ± SEM, n = 4. p = One-way ANOVA with Tukey’s post hoc analysis. **(F)** schematic of the two-way splint ligation approach. **(G)** RLUs were measured in lysates from HEK293T cells transfected with ligated and gel-purified NanoLuc mRNAs (n = 8) or the mRNA reporter produced by two-way-splint ligation (n = 4). Both constructs carried 2’-O-methylation at position A[+1] of the AUG start codon. Mean ± SEM. p = Two-tailed Student’s t-test.

To test the specificity of this system, we first transfected HEK293T cells with the ligated NanoLuc mRNAs containing a start codon, ligated NanoLuc mRNAs lacking a start codon, or unligated RNA pieces (Fig. 2C, Supplementary Fig. S2D). The ligated mRNAs containing the start codon within a Kozak sequence exhibited an ∼100-fold increase in NanoLuc activity compared to the unligated RNA pieces or the mRNAs lacking a start codon (Fig. 2C), demonstrating that ligation of a start codon is needed for translation of the mRNA reporters. To control for possible confounding artifacts, wherein a high proportion of unligated products affects translation, we transfected increasing concentrations of full-length NanoLuc mRNA with constant amounts of unligated RNA (Fig. 2D). Reciprocally, we transfected constant amounts of full-length NanoLuc mRNA with increasing concentrations of unligated RNAs (Supplementary Fig. S2E). In either case, NanoLuc activity was unaffected by the unligated RNAs, exhibiting linear behavior across a broad range of mRNA concentrations.

We next ligated unmodified or 2’-O-methylated Kozak sequences. Ligation efficiency was verified by RT-qPCR, yielding similar values across multiple replicates (Supplementary Fig. S2F and G). Equivalent amounts of ligated mRNAs were transfected into HEK293T cells, and translation efficiency was measured six hours after transfection through a luminescence assay. Remarkably, methylation of the 2’-OH group in the first position of the AUG codon significantly reduced NanoLuc activity to background levels (Fig. 2E). Significant luminescence reduction was also observed when 2’-O-methylation occurred on positions G[+3] and G[+4] (Fig. 2E). In contrast, no effect was observed in positions A[-3], C[-2], C[-1], and U[+2] (Fig. 2E). The inhibition of NanoLuc activity by A_m_[+1] remained after normalization of NanoLuc activity by transfected mRNA levels (Supplementary Fig. S2H).

To ensure that the NanoLuc signal was not influenced by unligated RNA fragments or potential artifacts from the three-way splint ligation, we used two complementary validation approaches (Supplementary Fig. S2I). First, we gel-purified full-length unmodified or A_m_[+1] mRNA reporters (Supplementary Fig. S2J and K). For this experiment, a longer 5’UTR was used to facilitate clear separation of the RNA fragments during gel purification (Supplementary Fig. S2I and J). Second, we carried out a two-way splint ligation using a short (27 nt) 5’ RNA fragment that was capped and 2’-O-methylated through chemical synthesis (Fig. 2F). The two-way splint ligation achieved nearly 100% ligation efficiency (Supplementary Fig. S2L and M).

Transfection of either gel-purified three-way ligated mRNA reporters or the two-way ligated counterpart produced results consistent with those obtained using unpurified mRNA reporters: the A_m_[+1] modification significantly inhibited NanoLuc activity to background levels (Fig. 2G). Although the two-way splint ligation approach provides high ligation and translation efficiency (Fig. 2G), chemically synthesized RNA has a 5’-OH end, which is unsuitable for enzymatic capping. Consequently, chemical capping is required, which substantially increases costs and limits the maximum length of RNAs that can be synthesized. Therefore, the two-way method is not cost-effective for systematically screening the site-specific effect of multiple RNA modifications in mRNA reporters. Taken together, our findings support the use of unpurified splint-ligated mRNA reporters to screen site-specific effects of 2’-O-methylation in translation regulation.

We next evaluated the reproducibility of our finding across different mRNA reporters and cell types. First, we transfected the ligated NanoLuc mRNAs into HeLa cells. Like in HEK293T cells, A_m_[+1] inhibited NanoLuc activity to background levels in HeLa cells (Supplementary Fig. S3A). G_m_[+3] and G_m_[+4] significantly reduced NanoLuc activity, albeit to a lesser extent than A_m_[+1] (Supplementary Fig. S3A). Second, we generated green fluorescent protein (GFP) mRNA reporters, each one containing 2’-O-methylation at a different position in the Kozak sequence (Supplementary Fig. S3B and C). Following transfection of the GFP mRNA reporters into HEK293T cells, protein expression was determined by flow cytometry. Consistently, the GFP fluorescence results mirror those of NanoLuc activity: A_m_[+1] inhibited GFP expression to baseline levels; G_m_[+3] and G_m_[+4] reduced protein expression, whereas A_m_[-3], C_m_ [-2], C_m_ [-1], and U_m_ [+2] had no effect on GFP expression (Fig. 3A and B).

**Figure 3.**
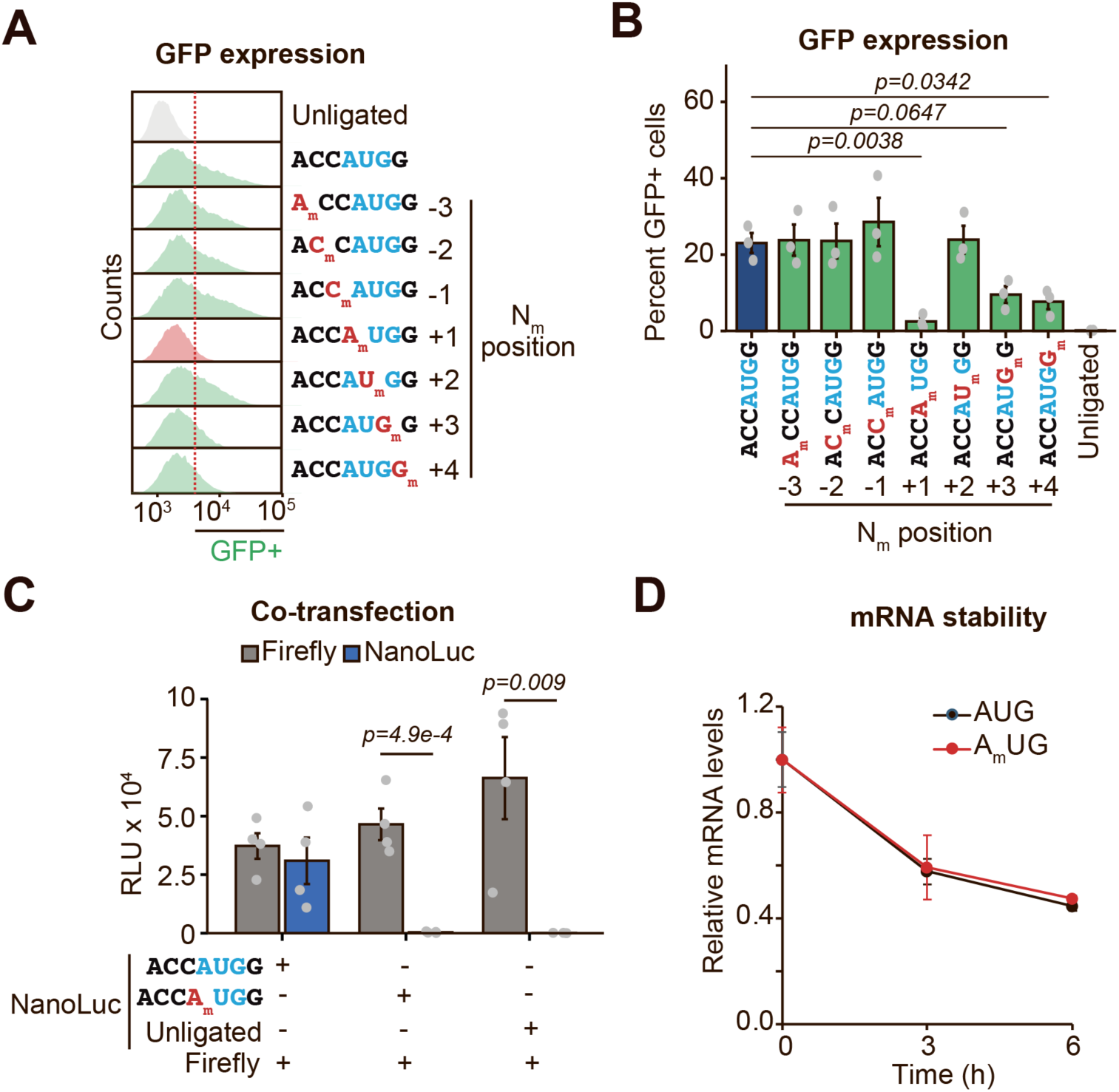
2’-O-methylation of A[+1] within start codons inhibits translation in a transcript-specific manner. **(A)** representative histograms of fluorescence intensity measured by flow cytometry in HEK293T cells transfected with ligated GFP mRNA reporters carrying 2’-O-methylation in different positions around the start codon. **(B)** quantification of data from (A). Mean ± SEM, n = 3. p = One-way ANOVA with Tukey’s post hoc analysis. **(C)** relative light units were measured in lysates from HEK293T cells co-transfected with unmodified mRNAs encoding Firefly luciferase and unmodified or A_m_[+1]-modified mRNAs encoding NanoLuc. Mean ± SEM, n = 4. p = Two-Tailed Student’s *t-test*. **(D)** NanoLuc mRNA levels were measured using RT-qPCR at different time intervals post-transfection. Primers were designed to detect only ligated mRNA. 18S rRNA was used as a normalizing control. Mean ± SEM, n = 3.

To guarantee that the inhibitory effect of A_m_[+1] on NanoLuc activity is transcript-specific and not a global perturbation of protein synthesis, we co-transfected unmodified or A_m_[+1]-modified NanoLuc mRNAs with unmodified Firefly luciferase mRNAs (Supplementary Fig. S3D). We observed that A_m_[+1] decreased NanoLuc activity to background levels without altering Firefly luciferase activity (Fig. 3C). In addition, evaluation of mRNA levels at different time intervals post-transfection showed no difference in mRNA stability between unmodified and 2’-O-methylated transcripts (Fig. 3D), indicating that A_m_[+1] inhibits translation only in 2’-O-methylated mRNAs.

In summary, consistent translation inhibition by A_m_[+1] was observed across unpurified and gel-purified three-way ligations, two-way ligations, three types of 5’UTRs (170 nt, 504 nt, and 27 nt), two reporters (NanoLuc, GFP), two cell types (HEK293T, HeLa), and was not attributable to mRNA instability. Since 2’-O-methylation eliminates the donor capacity of 2’-OH groups, these findings suggest that H-bonds between the 2’-OH group of nucleotide A[+1] and U1830 in 18S rRNA are relevant during translation initiation.

### 2’-O-methylation reduces translation from non-AUG start codons

We previously performed Ribo-seq in the presence of harringtonine (8), a translation inhibitor that allows the mapping of translation initiation sites transcriptome-wide (Supplementary Fig. S4A) (8). These experiments identified alternative translation initiation sites in ∼70% of cellular mRNAs (Supplementary Fig. S4B). Distribution of upstream start codons shows a higher prevalence of CUG, AUG, and GUG, with evidence for upstream initiation in all near-cognate start codons (Fig. 4A). Building on these observations, we sought to determine whether 2’-O-methylation affects translation initiation in the most prevalent near-cognate start codons: CUG and GUG. Constructs were produced by three-way splint ligation followed by gel purification (Fig. 4B and Supplementary Fig. S4C).

**Figure 4.**
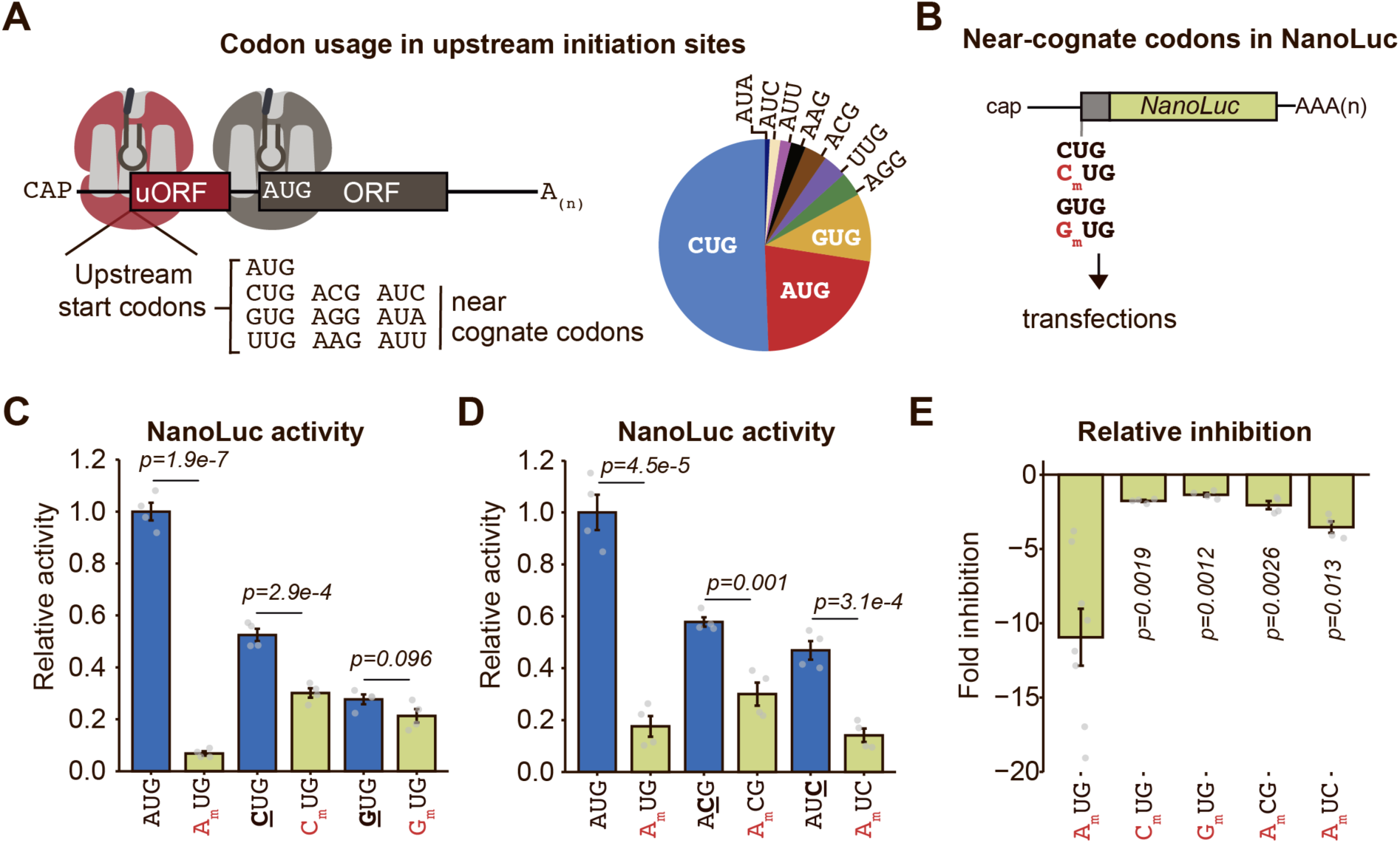
Inhibition of non-AUG translation by 2’-O-methylation. **(A)** distribution of upstream initiation codons obtained from HR-Ribo-seq (Arango et al., 2022) (8) **(B)** schematic of mRNA reporters with CUG and GUG initiation codons. **(C-D)** RLUs were measured in lysates from HEK293T cells transfected with gel-purified mRNA reporters carrying 2’-O-methylation in position N[+1] or left unmodified. RLUs were normalized relative to AUG. Mean ± SEM, n = 4. p = two-tailed *Student’s t-test*. **(E)** the relative inhibition effect from (C-D) was calculated as the fold change between the methylated and unmethylated conditions. p = One-way ANOVA with Tukey’s post hoc analysis.

Expectedly, the unmodified CUG and GUG start codons exhibited reduced NanoLuc activity compared to AUG (Fig. 4C). A_m_UG reduced NanoLuc activity by ∼11-fold, whereas C_m_UG significantly inhibited NanoLuc activity by only 1.8-fold (Fig. 4C). In contrast, a non-significant effect was observed for G_m_UG, although the NanoLuc levels for unmodified GUG were as low as those of C_m_UG (Fig. 4C). Given the more substantial effect of 2’-O-methylation in AUG codons, we sought to determine whether this observation is an adenosine-specific effect or a cognate codon-specific effect. Thus, we tested A_m_CG and A_m_UC, which retain adenosine at position [+1] but differ at positions [+2] and [+3], respectively (Supplementary Fig. S4D). Though A_m_CG and A_m_UC significantly inhibited NanoLuc activity, with a ∼2-fold decrease for A_m_CG and a ∼3.5-fold decrease for A_m_UC (Fig. 4D), the effect of 2’-O-methylation in all near-cognate codons was diminished compared to A_m_UG (Fig. 4E). These findings indicate that 2’-O-methylation exerts the most substantial effect in cognate AUG start codons.

Based on the modest inhibitory effect of 2’-O-methylation in near-cognate codons, we revisited cryo-EM structures of eukaryotic translation initiation complexes on near-cognate initiation. The only structure available was for the human 48S preinitiation complex on AUC, which shows an absence of the H-bond between A[+1] in mRNA and U1830 in 18S rRNA (Supplementary Fig. S4G).

Collectively, the presence of a putative hydrogen bond between the 2’OH group of A[+1] and 18S rRNA, the absence of this H-bond in near-cognate initiation, the strong inhibitory effect of 2’-O-methylation in A_m_UG, which disrupts the hydrogen bond capacity of the 2’-OH group, and the less significant effect of 2’-O-methylation in near-cognate codons, suggests that the interaction of 18S with A[+1] strengthens the optimality of Kozak sequences.

### 2’-O-methylation inhibits start codon recognition

Possible explanations for the inhibitory effect of 2’-O-methylation in translation initiation include: i) 2’-O-methylation prevents loading of the 40S subunit into mRNA; ii) 2’-O-methylation in position A[+1] stalls or induces disassociation of the 48S preinitiation complex before or after recognizing the start codon; iii) A_m_[+1] stalls or induces disassociation of the 80S initiation complex; or iv) 2’-O-methylation in position A[+1] makes the start codon non-optimal or unrecognizable by the preinitiation complex.

To begin testing these possibilities, we generated uORF-reporter systems in which an upstream uORFs is controlling the expression of NanoLuc (Supplementary Fig. S5A-C). We then incorporated 2’-O-methylation at the upstream AUG, leaving the downstream AUG unmodified. Expectedly, uORF inclusion decreased NanoLuc activity (Fig. 5A and B). Notably, we observed that 2’-O-methylation in the upstream AUG codons exhibited significantly higher NanoLuc activity compared to unmodified mRNA reporters, indicating that A_m_[+1] inhibits recognition of the upstream start codon and promotes downstream initiation (Fig. 5A and B). This observation suggests that 2’-O-methylation does not block the 40S subunit from engaging mRNAs and does not induce stalling/dissociation of the 48S or 80S complexes. Instead, the results of Fig. 5A and B suggest that A_m_[+1] hinders start codon recognition, allowing the 48S complex to initiate at a downstream start codon.

**Figure 5.**
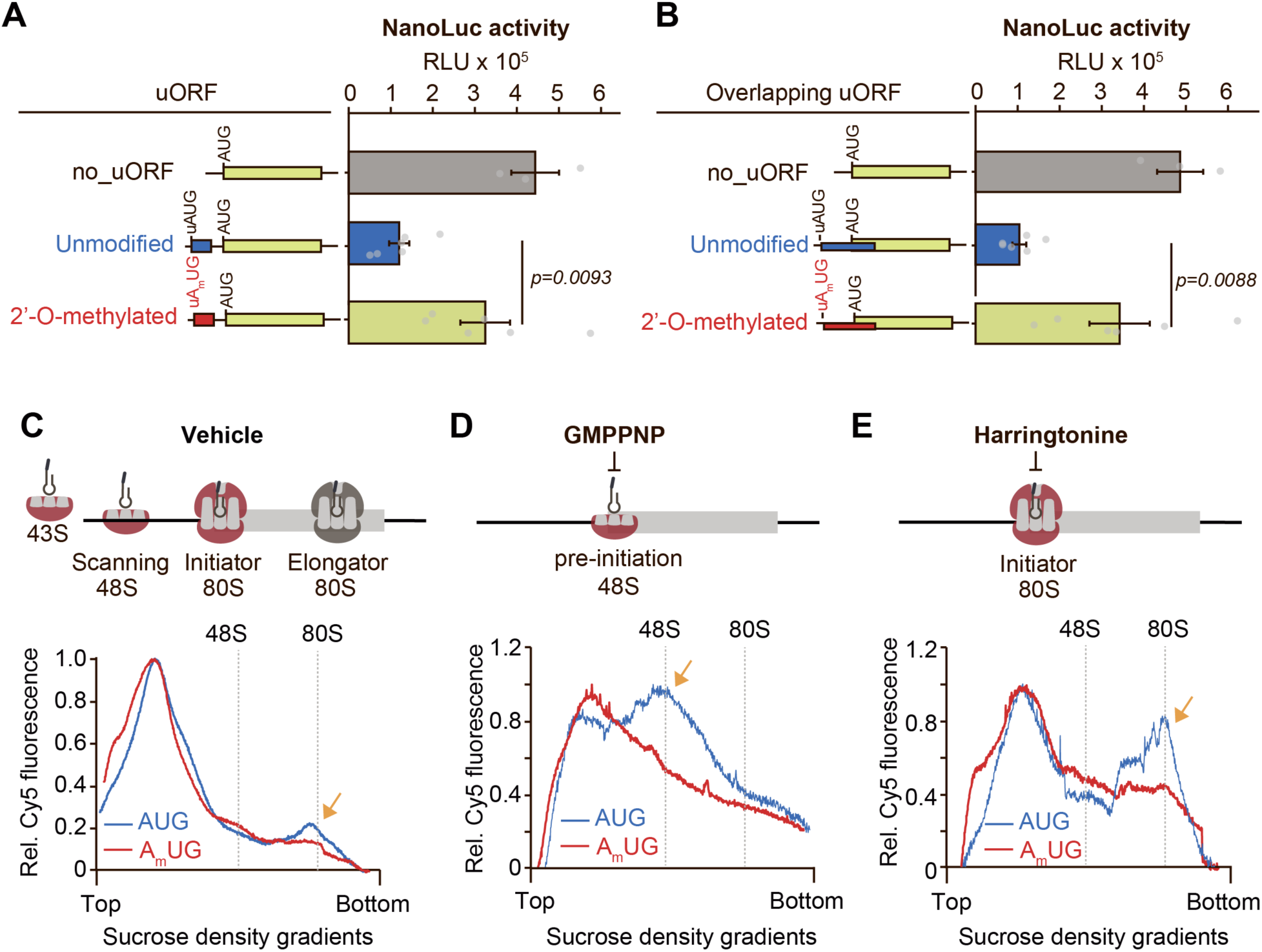
2’-O-methylation inhibits start codon recognition. **(A-B)** mRNA reporters carrying 2’-O-methylated or unmethylated uORFs were generated by two-way splint ligation and transfected into HEK293T cells. RLUs were measured six hours after transfection. Mean ± SEM, n = 4. p = two-tailed *Student’s t-test*. **(C-E),** *top:* schematic representation of translation initiation (C), 48S inhibition by GMPPNP (D), and 80S inhibition by harringtonine (E). *bottom:* Cy5-labeled synthetic mRNAs containing unmodified or A_m_UG-modified start codons were incubated with untreated reticulocyte lysates (C), treated with GMPPNP (D), or treated with harringtonine (E). Reactions were separated on a sucrose density gradient followed by fluorescence (Cy5) recording.

To validate these findings, we incubated reticulocyte lysates with a Cy5-labeled mRNA reporter in the presence of non-hydrolyzable GTP (GMPPNP) or harringtonine (HR) (Fig. 5C-E). Pre-treatment with GMPPNP stalls the 48S preinitiation complex after codon recognition (Fig. 5D). Relatedly, HR stalls the 80S initiation complex before proceeding into elongation (Fig. 5E). Reactions were separated on a 5-30% sucrose density gradient followed by fluorescence (Cy5) recording. As expected, treatment with GMPPNP increases the abundance of the 48S peak compared to vehicle control (Fig. 5D, blue lines), whereas harringtonine treatment increases the 80S peak in the Cy5-labeled unmodified mRNA reporter (Fig. 5E, blue line). In contrast, A_m_[+1] decreases the formation of both the 48S peak and 80S peaks in the presence of GMPPNP or HR, respectively (Fig. 5C-E, orange arrows).

Altogether, these findings support a model in which 2’-O-methylation at N[+1] reduces start codon recognition and favors downstream initiation.

### 2’-O-methylation is present in near-cognate codons within 5’UTRs

Since N_m_ naturally occurs within mRNAs (17), our findings raise the possibility that 2’-O-methylation is a regulatory mechanism of translation initiation in cellular mRNAs. Using Nanopore direct RNA sequencing (DRS), members of this team previously identified N_m_ sites in HeLa and C4-2 cell lines (Supplementary Fig. S6A) (17). We re-analyzed these Nanopore data sets to determine whether 2’-O-methylation is present in canonical AUG start codons. To increase confidence in the candidate N_m_ sites, only sites with an estimated modification ratio of more than 10% per position were included. Notably, no A_m_UG sites in canonical start codons were identified in either HeLa or C4-2 cell lines (Supplementary Fig. S6B).

We next extracted the N_m_ sites that mapped to 5’UTRs of any annotated Ensembl transcript and examined whether these sites were located in position N[+1] of an upstream AUG or an upstream near-cognate codon. This analysis identified 151 N_m_[+1] sites in HeLa cells and 181 sites in C4-2 cells (Supplementary Fig. S6B and Supplementary Tables S3 and S4). These sites were located in genes involved in biological pathways and molecular functions related to ribosome function and translation (Supplementary Fig. S6C). Notably, 65 sites (∼40%) were commonly observed in the 5’UTRs of HeLa and C4-2 cells, with only two A_m_UG sites (Supplementary Fig. S6D and Supplementary Table S5). The first A_m_UG site mapped to the ribosome biogenesis factor GNL3, although this site is polymorphic in the human population (rs1108842, A>C: ∼50%) (6). For instance, HeLa cells carry a CUG at this position (Supplementary Fig. S6E). The second AUG site mapped to the ribosomal protein RPS3A. However, this site is located within a region that could be part of the coding sequence or the 5’UTR, depending on the transcript isoform (Supplementary Fig. S6F). These observations indicate that A_m_UG is either absent or present at very low stoichiometry in HeLa and C4-2 cells. Nevertheless, N_m_ sites were identified in near-cognate codons within 5’UTRs (Supplementary Fig. S6B). CUG had the highest prevalence, followed by AUC (Supplementary Fig. S6B), with C_m_UG and A_m_UC sites significantly enriched relative to the distribution of these codons in 5’UTRs (Supplementary Fig. S6G).

To validate the presence of N_m_[+1] in near-cognate codons, we employed <u>R</u>everse <u>T</u>ranscription at <u>L</u>ow dNTPs (RTL-P) (25). This method leverages the property of 2’-O-methylation to induce reverse transcriptase stops (RT-STOP) at low concentrations of dNTPs (Fig. 6A), resulting in a reduction in amplicon density after PCR amplification of the RT product (Supplementary Fig. S7A, red arrowhead). While there are more than eleven human 2’O-methyltransferase enzymes (26), fibrillarin (FBL) is consistently identified as the predominant enzyme responsible for introducing N_m_ at internal sites in mRNA (17–20,27). Thus, we performed RTL-P in total RNA isolated from HeLa cells transfected with two different siRNAs against Fibrillarin (FBL) or a siRNA control (Fig. 6B). Candidate sites were selected based on the highest predicted modification ratio from the Nanopore-DRS approach in HeLa cells (Supplementary Table S6). Of the 12 candidate sites tested, 10 exhibited a substantial loss in amplicon density under limited dNTP conditions, with relative methylation values ranging from 30% to 99% in the validated sites (Fig. 6C-F and Supplementary Fig. S7A-B). The RTL-P signature was sensitive to FBL knockdown (Fig. 6G-I and Supplementary Fig. S7B), validating the presence of 2’-O-methylation sites in near-cognate codons within human 5’UTRs.

**Figure 6.**
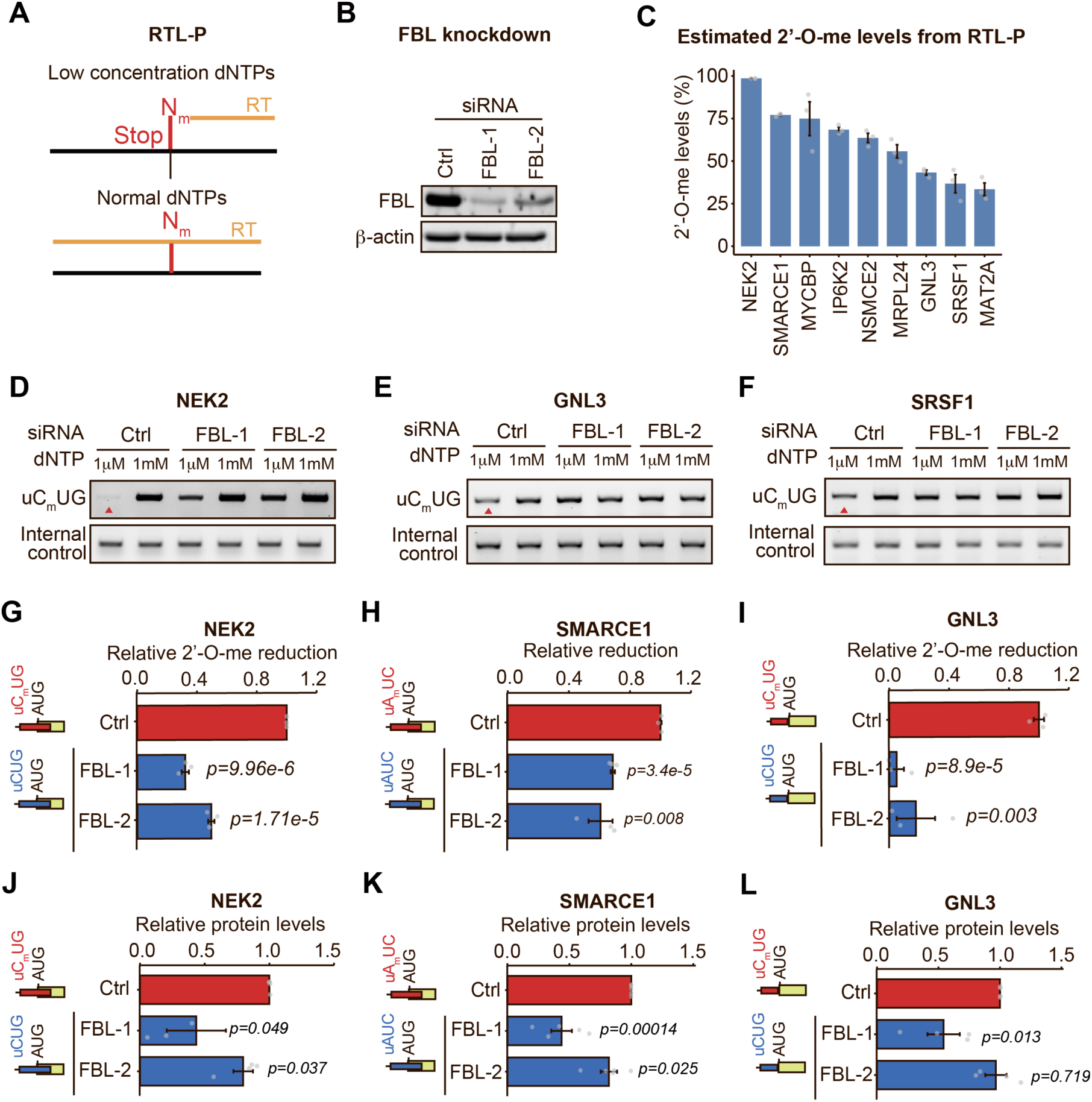
Identification of 2’-O-methylation in start codons of cellular mRNAs. **(A)** schematic of RTL-P. Reverse transcription was performed in the presence of an optimal dNTP concentration (1 mM) or a suboptimal dNTP concentration (1μM) that induces a premature RT-STOP at 2’-O-methylation sites. A pair of primers surrounding the 2’-O-methylation is used during PCR amplification. In this manner, the amplicon signal is an indicator of RT-STOP efficiency and N_m_ stoichiometry. Another pair of primers, located downstream of the 2’-O-methylation site, serves as an internal control for mRNA levels. **(B)** Western blot validation of FBL knockdown using two different siRNAs in HeLa cells. **(C)** quantification of the relative N_m_ ratio from the gels shown in Fig. 6D-F and Supplementary Fig. 7A. Mean ± SEM, n = 3. **(D-F)** representative agarose gels of amplicon density for the 2’-O-methylated site (top) and the loading control (bottom). Arrowheads indicate the reduction in amplicon signal when reverse transcription is performed at low dNTP concentration in the presence of FBL. **(G-I)** quantification of the relative N_m_ ratio dependent on FBL from the agarose gels shown in (D-F). Mean ± SEM, n = 3. p = two-tailed *Student’s t-test*. (**J-L),** quantification of the relative protein levels dependent on FBL knockdown from the gels shown in Supplementary Fig. S7D. Mean ± SEM, n = 4-5. p = two-tailed *Student’s t-test*.

To evaluate whether 2’-O-methylation in upstream near-cognate codons affects translation, we analyzed publicly available Ribo-seq data from HeLa cells transduced with an inducible shRNA against FBL. Translation efficiency (TE) was calculated for genes with N_m_[+1] in upstream near-cognate codons compared to genes with no 2’-O-methylation in the 5’UTR. The results suggest that loss of N_m_[+1] in upstream near-cognate codons, due to FBL knockdown, significantly decreases translation efficiency compared to genes with no 2’-O-methylation in the 5’UTR (Supplementary Fig. S7C and Supplementary Table S7). These findings are consistent with the uORF reporter assays, where reporters with unmodified uORFs exhibited significantly less NanoLuc activity compared to N_m_[+1] in upstream start codons (Fig. 5A and B).

Lastly, we validated the effect of upstream 2’-O-methylation on canonical protein expression using Western blots. We chose NEK2, GNL3, and SRSF1, which had N_m_[+1] levels of ∼99%, ∼75%, ∼45%, and ∼30%, respectively, in upstream CUG or AUC codons (Fig. 6C). These sites could generate uORFs with inhibitory potential (Fig. 6G-I and Supplementary Fig. 7B, schematic), in which loss of 2’-O-methylation, due to FBL knockdown, would increase upstream CUG or AUC usage, leading to decreased canonical protein expression. NEK2 and SMARCE1 showed a consistent decrease in canonical protein expression among the two different siRNAs, with a more substantial effect in the siRNA-FBL-1 condition (Fig. 6J-K and Supplementary Fig. S7D). Notably, the siRNA-FBL-1 condition exhibited a more pronounced FBL knockdown (Fig. 6B). The consequence of FBL knockdown on GNL3 expression was weaker, with significant effects observed only in the siRNA-FBL-1 condition (Fig. 6L and Supplementary Fig. S7D). In contrast, SRSF1 protein levels were unchanged upon FBL knockdown (Supplementary Fig. S7D-E). These findings indicate that endogenous 2’-O-methylation at upstream CUG or AUC codons significantly affects downstream protein expression at high 2’-O-methylation levels, with a modest or imperceptible effect at lower levels. Notably, the predicted stoichiometries of 2’-O-methylation in upstream near-cognate codons from Nanopore DRS show a median of ∼20% (Supplementary Fig. S7F), explaining the modest, yet significant, effect of N_m_[+1] loss on translation efficiency transcriptome-wide in HeLa cells (Supplementary Fig. S7C).

Collectively, these findings confirmed N_m_[+1] in upstream near-cognates in HeLa cells, where it regulates the expression of downstream canonical proteins.

## DISCUSSION

The results presented in this study suggest that 2’-O-methylation inhibits translation initiation by interfering with start codon recognition, thereby allowing the scanning ribosome to initiate at downstream start codons (graphical abstract). This model is supported by the presence of a hydrogen bond between the 2’OH bond of AUG in mRNA and 18S rRNA (Fig. 1C), the absence of this H-bond in near-cognate initiation (Supplementary Fig. S4E), the strong inhibitory effect of 2’-O-methylation in A_m_UG (Fig. 2 and 3), a modification that disrupts the hydrogen bond capacity of 2’-OH groups (28), and the less significant effect of 2’-O-methylation in near-cognate codons (Fig. 4C-E). In addition, loss of N_m_[+1] at upstream start codons decreased downstream translation (Fig. 5A-B, Fig. 6J-K, and Supplementary Fig. S7C) and reduced GMPPNP- and HR-trapped complexes (Fig. 5D-E), consistent with reduced start codon recognition at upstream methylated sites.

Chemically, the 2’-OH group in RNA can act as both a donor and acceptor in H-bonds. Methylation replaces the hydrogen atom of the 2’-OH with a -CH_3_ group, effectively eliminating its donor capability. Since the 2’-OH group of the A[+1] position can act as a donor in an H-bond with U1830, either with the phosphate (PO_4_^3-^) backbone or the keto (C=O) group at C2, 2’-O-methylation is likely to block such interaction. We speculate that the A[+1]^mRNA^:U1830^18S^ interaction helps 18S rRNA monitoring for proper pairing and orientation of the codon:anticodon interaction. For instance, the movement of the 40S subunit along the 5′ UTR is characterized by a conformational intermediate between the closed state, in which the tRNA_i_^met^ is fully inserted into the P site, trying to make an interaction with the mRNA codon, and the open state, in which the anticodon in tRNA_i_^met^ is distant from the mRNA (23,29). This conformational intermediate enables the preinitiation complex to proofread accommodation of optimal start codons (9,22,29–34). We speculate that blocking the A[+1]^mRNA^:U1830^18S^ interaction by 2’-O-methylation precludes the ribosome from proofreading start codon recognition. Alternatively, 2’-O-methylation can alter the local structure of mRNA (35), thereby preventing it from properly pairing with the anticodon base. Since 2’-O-methylation inhibits start codon recognition and promotes downstream initiation (Fig. 5), it likely promotes leaky scanning, allowing the 43S complex to continue scanning until it encounters a downstream start codon. However, our findings do not rule out the possibility that 2’-O-methylation promotes reinitiation, rather than leaky scanning, or introduces internal ribosome entry sites that enhance downstream initiation.

The inhibitory effect of 2’-O-methylation is less pronounced in near-cognate codons, likely because they lack the proper pairing and orientation of the cognate codons. However, we evaluated only the Kozak context most optimal for AUG initiation (ACC<u>AUG</u>G). We prioritized CUG, GUG, AUC, and ACG based on HR-Ribo-seq prevalence (Fig. 4A), the A[+1] position, and the prevalence of these codons in HeLa 5 “UTRs (Supplementary Fig. S6B), but focused on a single Kozak context. Additional Kozak variants, near-cognate codons, and structured 5’UTRs that favor non-AUG initiation remain to be tested to accurately rank the effects of 2’-O-methylation in near-cognate initiation.

Notably, 2’-O-methylation of the start codon does not impair mRNA stability or globally affect protein synthesis (Fig. 3C and D), indicating that its effect on translation initiation is transcript-specific. The effect of 2’-O-methylation in translation initiation is also site-specific. 2’-O-methylation of position N[-3], N[-2], N[-1], and N[+2] had no effect on protein output (Fig. 2E and 3B). An unexpected finding was the lack of effect at position N[-1], despite our previous report of a significant role for the C[-1]^mRNA^:t^6^A^tRNA^ interaction in stabilizing the initiation complex (8). However, 2’-O-methylation does not alter the acceptor capacity of the C-O-CH3 group, and thus, the C[-1]^mRNA^:t^6^A^tRNA^ hydrogen bonds may still occur with N_m_[-1]. 2’-O-methylation in positions N[+3] and N[+4] reduced protein output, albeit to a much lesser extent than N[+1] (Fig. 2E and 3B). Alike N[-1], the 2’-OH group in N[+3] is an acceptor (Supplementary Table S2), rather than a donor, and thus, 2’-O-methylation would not disrupt its hydrogen bond. The effect of N_m_[+4] on protein synthesis could be mediated through impairing elongation as the ribosome moves into the second and third codons. In support of the latter, previous studies indicated that 2’-O-methylation represses the translocation step during translation elongation by precisely inhibiting the interaction of mRNA with the monitoring bases of rRNA during the proofreading step (27,28).

It is well established that 2′-O-methylation is highly abundant in noncoding RNAs, including rRNA and tRNA, and occurs in the 5′ cap of all mRNAs in human transcriptomes (26). More recently, the presence of N_m_ at internal mRNA has been reported in multiple studies (17–20,36–38). By re-analyzing published Nanopore DRS datasets (17), we report the absence of 2’-O-methylation in position A[+1] of canonical and upstream AUGs in HeLa and C4-2 cells under normal laboratory growth conditions. However, our analysis does not conclusively rule out the possibility that AUG codons are methylated in human transcriptomes. A more comprehensive study using additional mapping methods such as RiboMeth-seq (39), N_m_-seq (20), RibOxi-seq (37), NJU-seq (19), N_m_-mut-seq (18), or 2OMe-seq (36) in multiple cell types and tissues across different conditions would help determine whether A_m_UG codons exist in human transcriptomes.

We validated the presence of N_m_ sites in near-cognate codons within the 5’UTRs of HeLa mRNAs (Fig. 6 and Supplementary Fig. S7) and observed that loss of upstream N_m_, due to FBL knockdown, reduces translation of downstream proteins, especially at high-stoichiometry sites (Fig. 6J-K and Supplementary Fig. S7C). These observations suggest that cells can leverage 2’-O-methylation to control non-AUG translation. However, FBL also catalyzes 2’-O-methylation of ribosomal RNA, other noncoding RNAs, and the coding sequences of mRNAs, exerting pleiotropic effects that could confound or mask the effect of FBL knockdown at low stoichiometric sites in 5’UTRs. Moreover, our findings do not preclude the possibility that near-cognate 2’-O-methylation plays a role under stress conditions when canonical cap-dependent translation is repressed, allowing non-AUG translation to serve as a stress response mechanism. In this manner, cells could block unwanted non-canonical initiation sites through 2’-O-methylation under specific cellular conditions.

The finding that 2’-O-methylation of upstream start codons inhibits uORF usage while enhancing canonical protein expression in transcript-specific manners has a significant impact on RNA therapeutics. Aberrant translation initiation has been reported in numerous diseases, with hundreds of pathogenic genetic variants in 5’UTRs linked to the introduction or destruction of uORFs (5,40–43). These pathological conditions would benefit from strategies that inhibit aberrant initiation sites while enhancing expression of the canonical ORF. Since 2’-O-methylation can be targeted to specific positions within transcripts (44–46), our findings open the door to investigating approaches that can guide 2’-O-methylation for the targeted inhibition of pathogenic translation initiation in human disease. Moreover, 2’-O-methylation could be incorporated into the 5’UTRs of synthetic mRNA medicines to enhance protein expression.

In sum, we identified a chemical modification of mRNA, 2’-O-Methylation, that unambiguously inhibits start codon recognition in a position-specific manner, a finding that could be leveraged to inhibit aberrant uORF expression or enhance downstream ORF expression for RNA therapeutics.

## ACKNOWLEDGMENTS

We thank the Flow Cytometry Core Facility at the Robert H. Lurie Comprehensive Cancer Center for flow cytometry services, and the Quest High-Performance Computing Facility at Northwestern University for computational resources. We thank all members of the Arango lab for insightful discussions.

## AUTHOR CONTRIBUTIONS

Adam Suh: Investigation, Methodology, Validation, Formal Analysis, Writing – review & editing. Stephanie Mou: Investigation, Methodology, Validation, Formal Analysis, Writing – review & editing. Emmely A. Patrasso: Investigation, Methodology, Validation, Formal Analysis, Writing – review & editing. Hannah Serio: Formal analysis, Data curation, Visualization, Writing – review & editing. Kevin Vasquez: Investigation. Rui Wang: Investigation. Smriti Sangwan: Formal Analysis, Visualization. Yang Yi: Investigation, Validation, Formal Analysis. Daniel Arango: Conceptualization, Data curation, Formal analysis, Funding acquisition, Methodology, Project administration, Resources, Supervision, Visualization, Writing – original draft, Writing – review & editing.

## SUPPLEMENTARY DATA

**Supplementary Figure S1.**
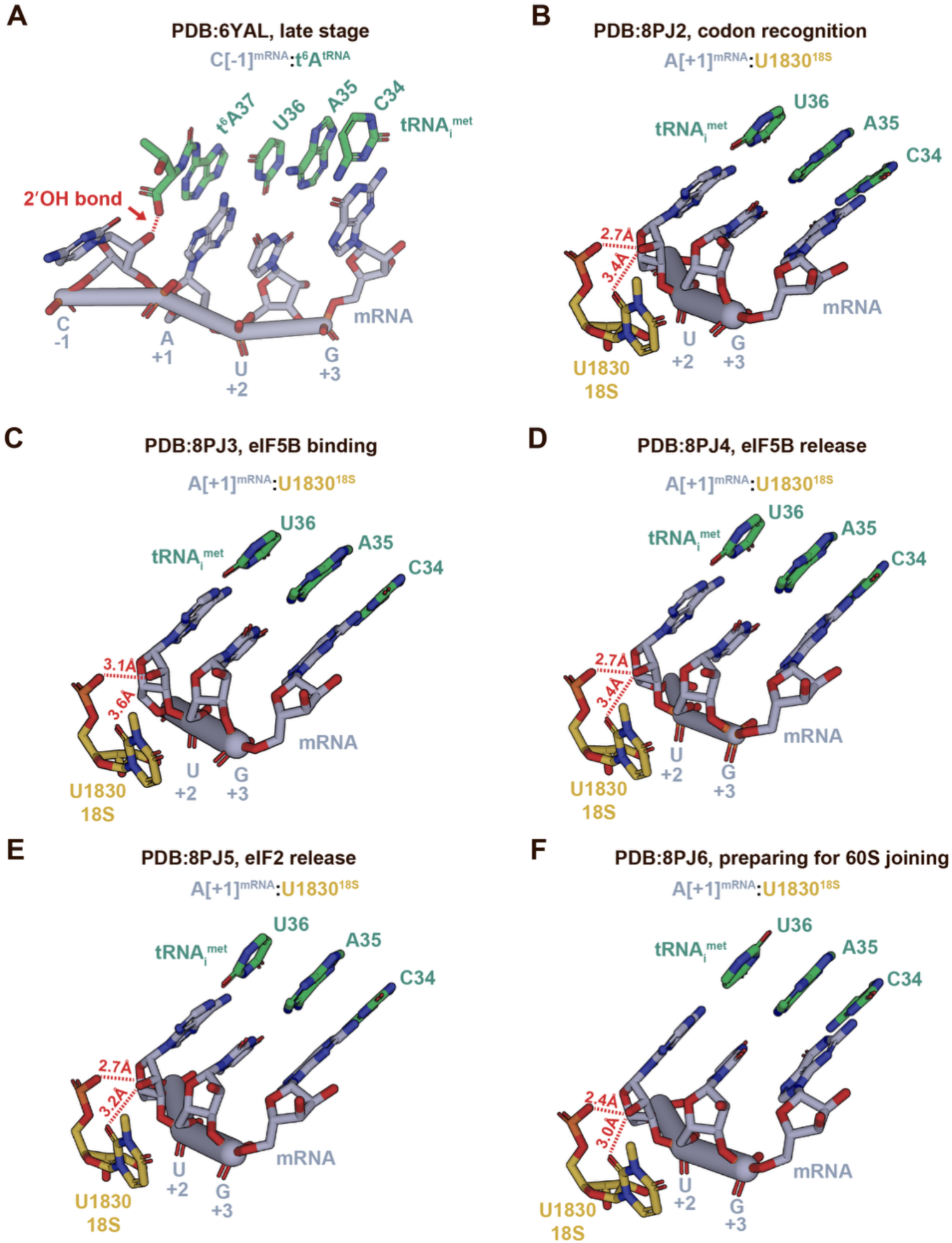
H-bonds between the 2’-OH groups of mRNA nucleotides with tRNA_i_^met^ and 18S rRNA. **(A)** Cryo-EM visualization of late-stage rabbit 48S preinitiation complexes (PDB:6YAL, Simonetti et al., 2022) (9) centered around the interaction of the 2’-OH group in nucleotide C[-1]^mRNA^ with t^6^A in tRNA_i_^met^. This interaction was reproduced in rabbit 80S initiator ribosomes in complex with Harringtonine (Fig. 1B). **(B-F),** Cryo-EM visualization of 48S preinitiation complexes (Petrychenko et al., 2025) (22) centered around the interaction of the 2’-OH group in nucleotide A[+1]^mRNA^ with U1830 in 18S rRNA across different stages of 48S preinitiation complexes, including the stage of codon recognition and GTP hydrolysis (B, PDB: 8PJ2), binding of eIF5B (C, PDB: 8PJ3), followed by sequential release of eIF5B (D, PDB: 8PJ4) and eIF2 (E, PDB: 8PJ5), and the late-stage readiness for 60S subunit joining (F, PDB: 8PJ6). All publicly available structures were downloaded from the PDB and visualized in PyMOL.

**Supplementary Figure S2.**
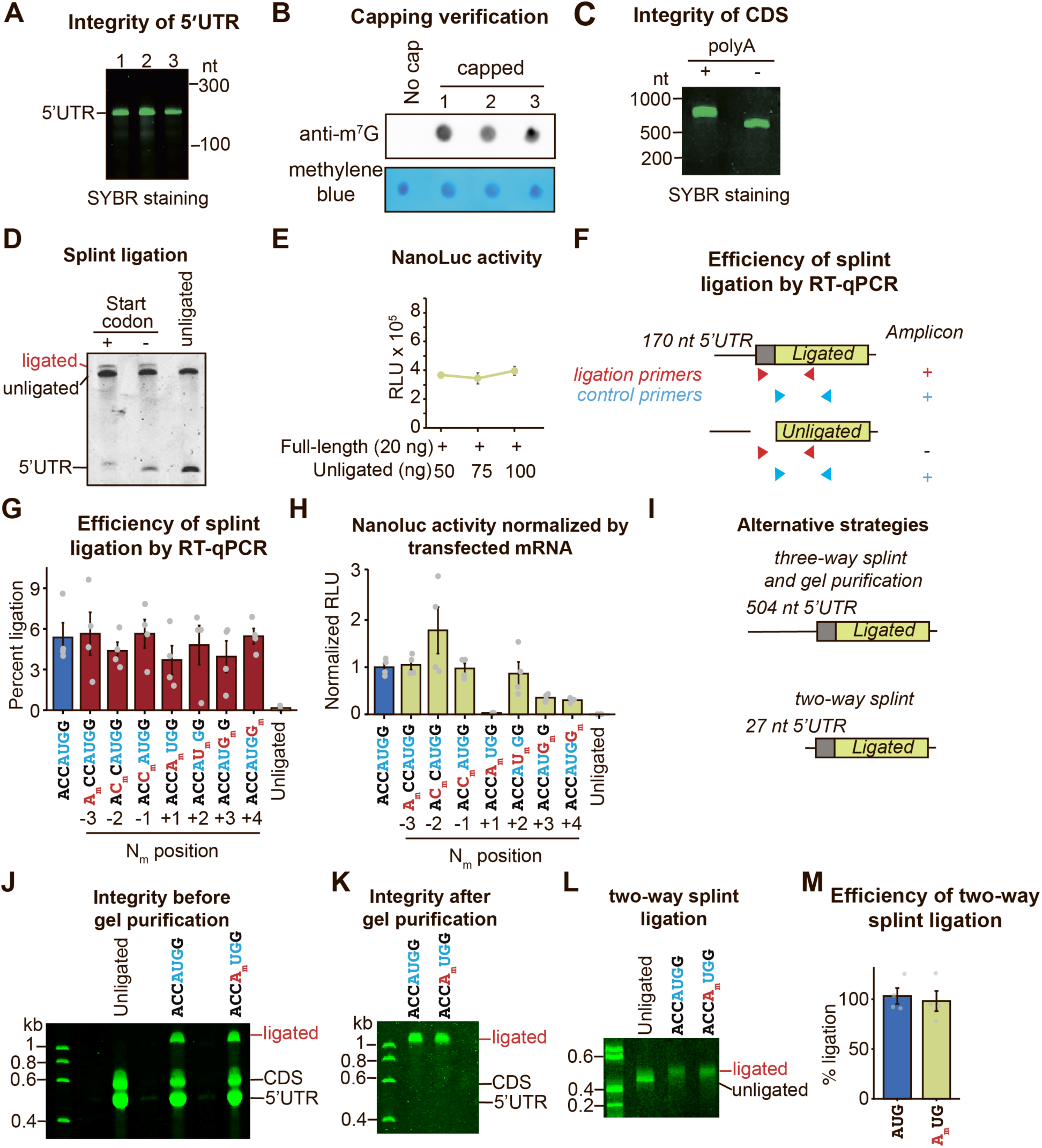
Quality controls for the construction of NanoLuc mRNA reporters. **(A)** the 5’ end of the mRNA reporters lacking a start codon was transcribed and capped *in vitro*. The integrity of the RNA was verified by polyacrylamide gel electrophoresis (PAGE) and SYBR gold staining. Three independent reactions are shown in the gel. **(B)** successful capping of the *in vitro* transcribed 5’ ends was verified by dot blot using anti-m^7^G antibodies. Methylene blue staining was used as a loading control. Three independent capping reactions are shown in the membrane. **(C)** the 3’ end of the mRNA reporters, containing the ORF and the 3’UTR but lacking a start codon, was transcribed and polyadenylated *in vitro*. The integrity of the RNA and the efficiency of polyadenylation were verified using PAGE and SYBR gold staining. **(D)** the ligation of NanoLuc mRNA reporters was verified using PAGE and SYBR gold staining. Ligated mRNA shows a shift in molecular weight compared to unligated mRNA. **(E)** relative light units were measured in lysates from HEK293T cells co-transfected with increasing amounts of unligated RNA pieces and constant amounts (20 ng) of full-length NanoLuc mRNA. Mean ± SEM, n = 3. The line plot shows stable NanoLuc activity across different amounts of unligated RNA. **(F-G)** ligation efficiency was verified by RT-qPCR. Two primer sets were used (Supplementary Table S1). The first set specifically recognizes the ligated mRNA, while the second set binds the NanoLuc ORF to obtain total mRNA levels. The efficiency of ligation was calculated as: (ligated mRNA)/(total RNA)*100. **(H)** NanoLuc activity was normalized to the amount of transfected mRNA, which was quantified by RT-qPCR. Mean ± SEM, n = 3. **(I)** schematic of different constructs used in three-way and two-way splint ligations. **(J)** the ligation of NanoLuc mRNA reporters was verified using PAGE and SYBR gold staining. Ligated mRNA shows a shift in molecular weight compared to unligated mRNA. **(K)** the integrity of the gel-purified mRNA reporters was verified using PAGE and SYBR gold staining. **(L)** the ligation of NanoLuc mRNA reporters was verified using PAGE and SYBR gold staining. Ligated mRNA shows a slight shift corresponding to a 40 nt difference in molecular weight compared to unligated mRNA. **(M)** The efficiency of the two-way ligation was verified by RT-qPCR. Mean ± SEM, n = 4.

**Supplementary Figure S3.**
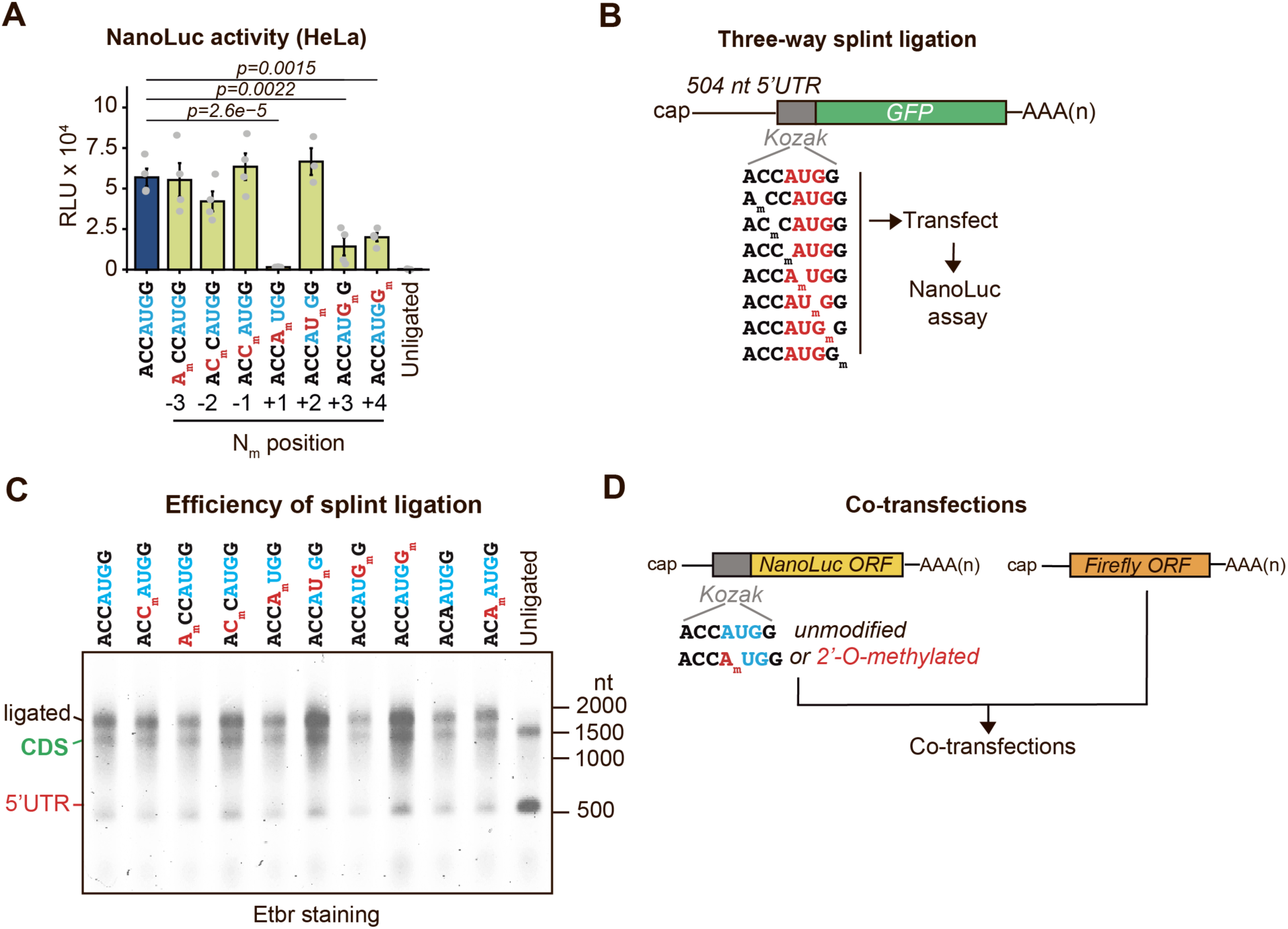
Quality controls for the construction of GFP mRNA reporters. **(A)** relative light units were measured in lysates from HeLa cells transfected with ligated NanoLuc mRNAs carrying 2’-O-methylation in different positions around the start codon. Unmethylated (blue) and unligated mRNAs were used as controls. Mean ± SEM, n = 4. p = One-way ANOVA with Tukey’s post hoc analysis. **(B)** schematic of the GFP mRNA reporters. **(C)** denaturing agarose gel electrophoresis, indicating the ligation efficiencies for the GFP mRNA reporters. These RNA fragments are larger than those used for NanoLuc reporters and do not resolve well in PAGE gels. Thus, agarose gels are better suited for this separation. **(D)** schematic indicating the co-transfection strategy of the full-length firefly luciferase as a control compared to the splint ligated NanoLuc mRNA reporter.

**Supplementary Figure S4.**
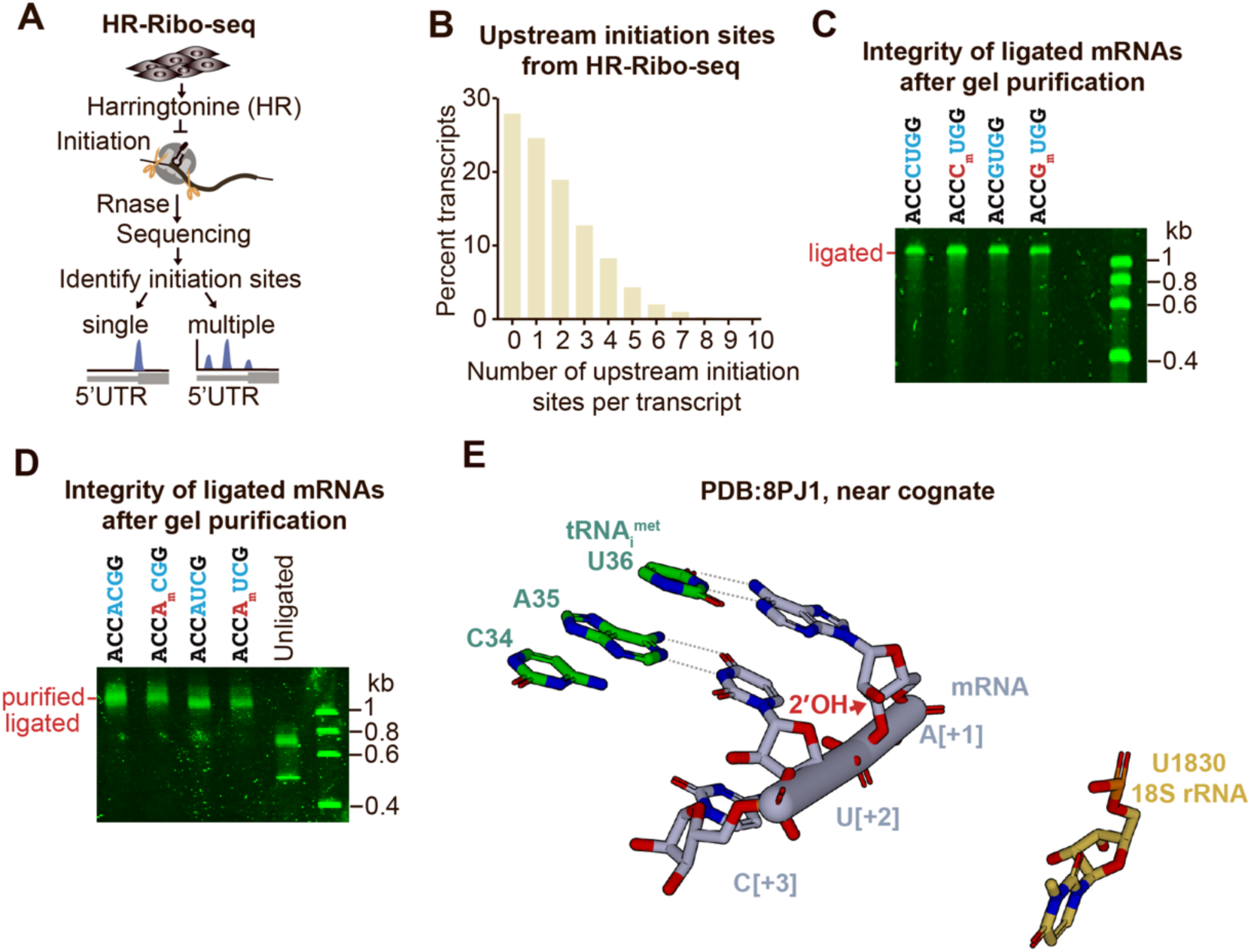
Quality controls for the construction of mRNA reporters with near-cognate initiation codons. **(A)** To determine the upstream near-cognate or AUG codons used for initiation, we previously performed HR-Ribo-seq in HeLa cells (8). In this method, the inhibitor Harringtonine (HR) stalls the initiator ribosome while letting the elongator ribosomes slide off of mRNA. Subsequent RNAse footprinting and isolation of ribosome-protected fragments (RPFs) allow the identification of initiation sites transcriptome-wide. **(B)** HR-Ribo-seq indicates that ∼70% of transcripts initiate translation from at least one upstream site in wildtype HeLa cells. **(C-D)** the integrity of ligated NanoLuc mRNAs carrying near-cognate codons was verified using PAGE and SYBR gold staining. **(E)** Cryo-EM visualization of 48S preinitiation complexes (Petrychenko et al., 2025) (22) centered around nucleotide A[+1]^mRNA^ from an AUC start codon (PDB: 8PJ1). The distance between U1830 in rRNA and A[+1] in mRNA (10.5Å) does not support the formation of an H-bond. It is important to note that AUC is in different contexts in the cryo-EM structure (AGG<u>AUC</u>C) than our mRNA reporter system (ACC<u>AUC</u>G).

**Supplementary Figure S5.**
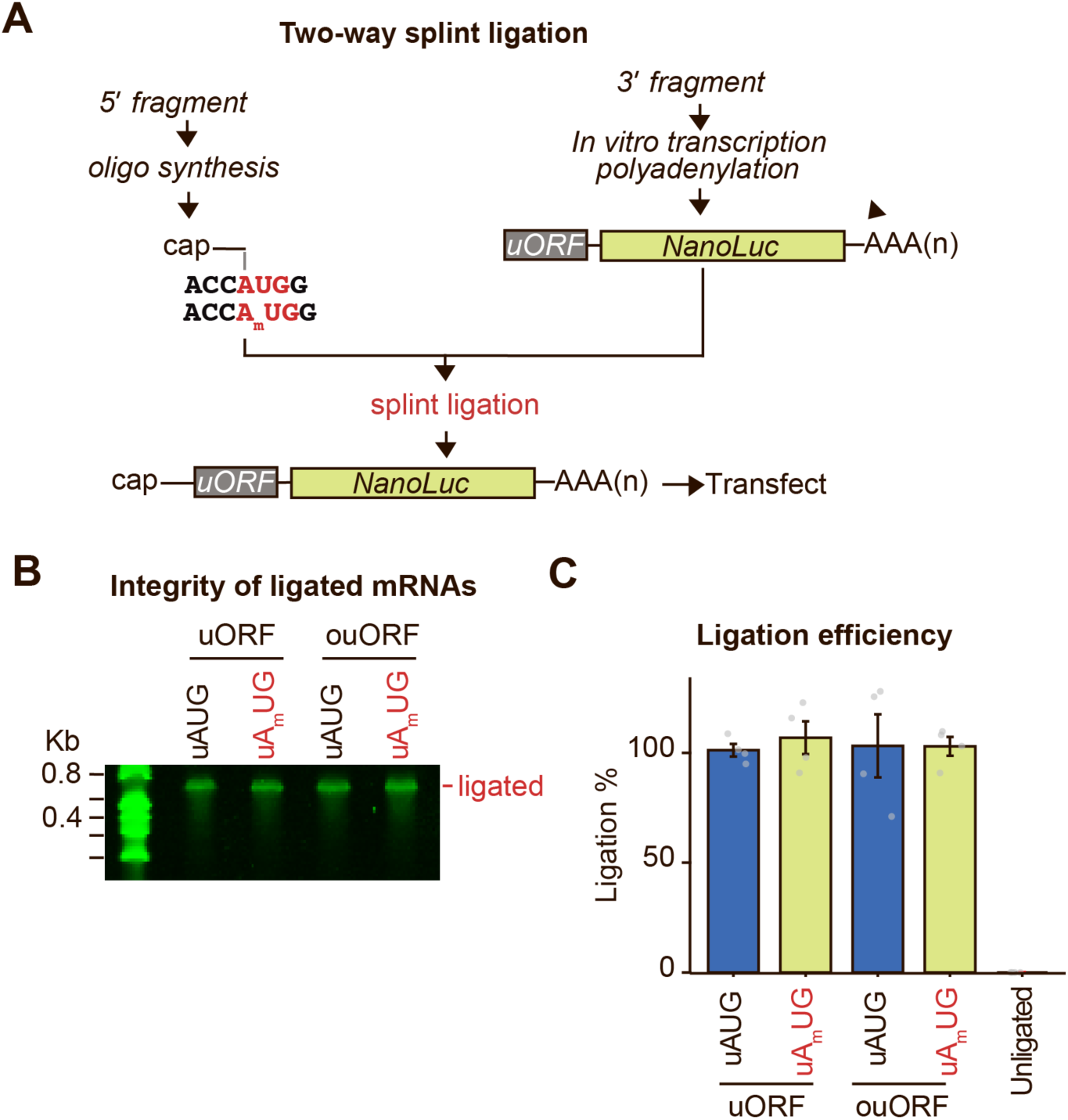
Quality controls for the construction of uORF reporters. **(A)** schematic of the two-way splint ligation strategy to generate mRNA reporters with 2’-O-methylated or unmethylated uORFs. **(B)** the integrity of ligated NanoLuc mRNAs was verified using PAGE and SYBR gold staining. **(C)** ligation efficiency was verified by RT-qPCR. Mean ± SEM, n = 4.

**Supplementary Figure S6.**
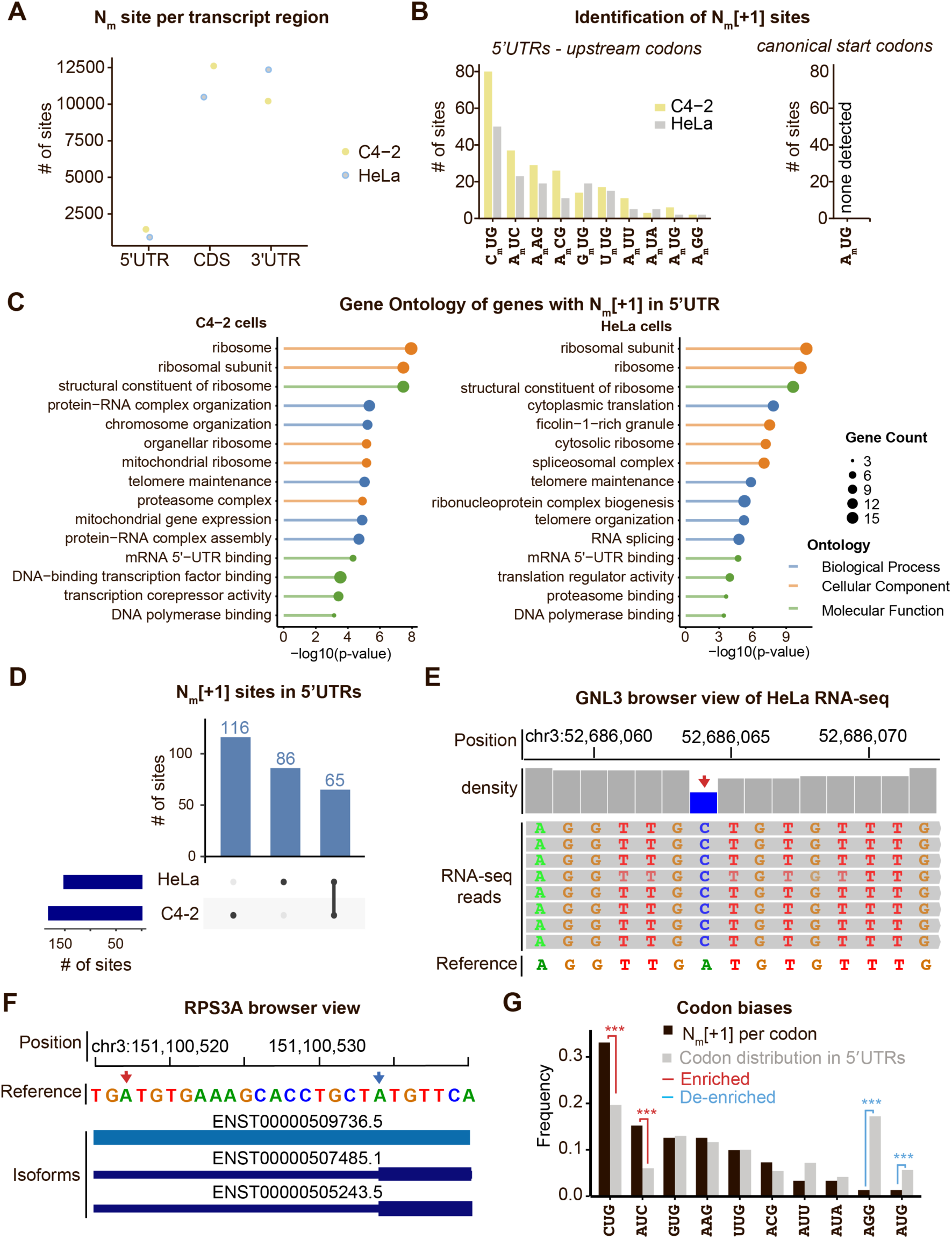
Distribution of 2’-O-methylation sites in human mRNAs. **(A)** 2’-O-methylation maps in HeLa and C4-2 cells were compiled and binned by transcript location (5’UTR, CDS, or 3’UTR)**. (B)** the genomic coordinate of 2’-O-methylation sites was intersected with the coordinates for canonical AUG, AUG within 5’UTRs, and near-cognate start codons within 5’UTRs of human transcriptomes. N_m_[+1] sites were binned by codon identity. **(C)** gene ontology analysis of all genes from (B). **(D)** an upset plot indicating the overlap of the sites described in (B) between HeLa and C4-2 cells. **(E)** browser view of RNA-seq reads in HeLa cells indicates an A>C SNP at an annotated upstream AUG codon. **(F)** browser view of different transcript isoforms in RPS3A indicates that the annotated AUG codon is located in the 5’UTR of some isoforms but in the coding sequence of other isoforms. Red arrow indicates the 2’-O-methylation sites. Blue arrow indicates the canonical start codon. **(G)** the fraction of each N_m_[+1]-methylated codon was compared to the distribution of AUG and all near-cognate codons in the 5’UTR of human mRNAs. p = chi-squared test for proportions. *** p < 0.0001.

**Supplementary Figure S7.**
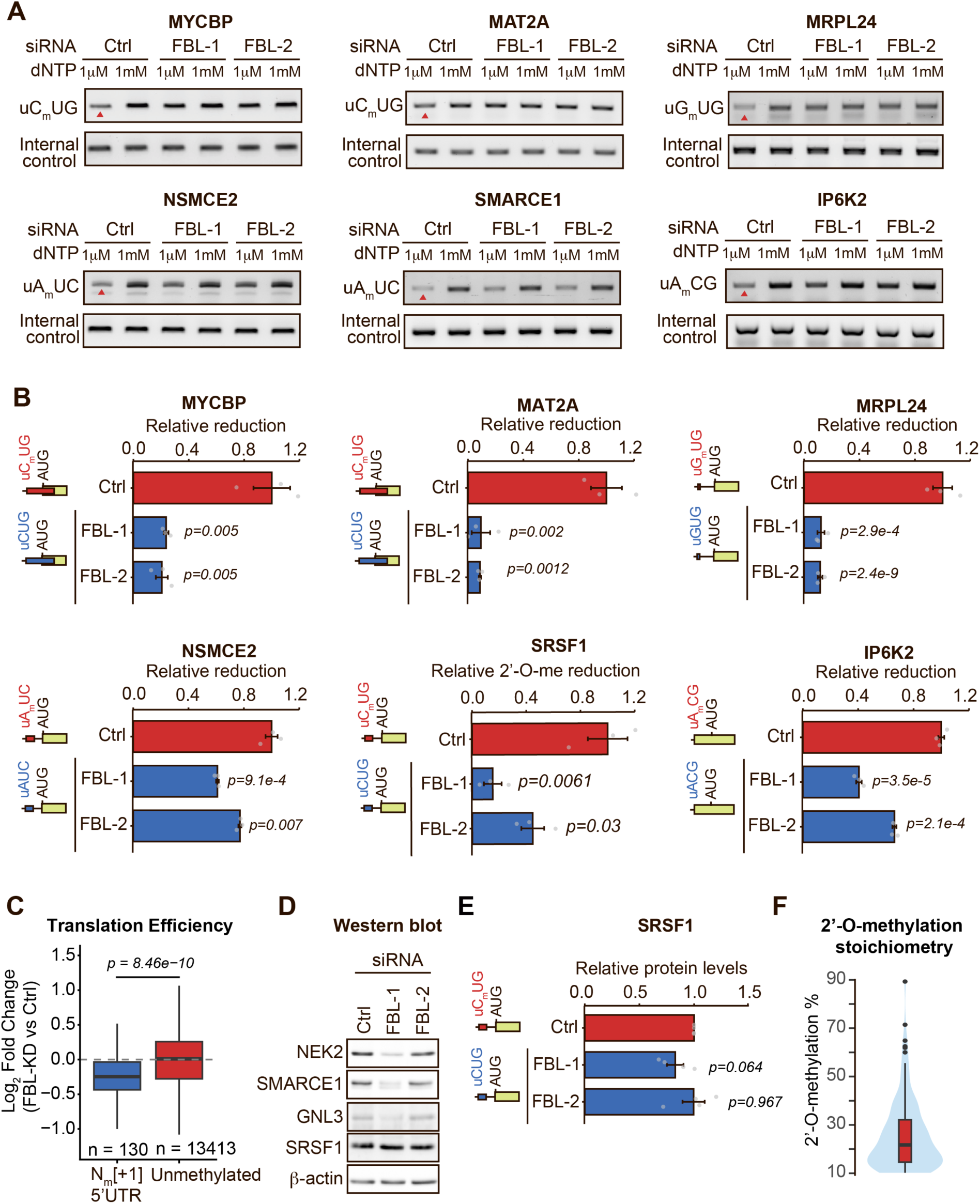
Validation of 2’-O-methylation sites in near-cognate codons. **(A)** representative agarose gels of amplicon density for the 2’-O-methylated site (top) and the loading control (bottom). Arrowheads indicate the reduction in amplicon signal when reverse transcription is performed at low dNTP concentration in the presence of FBL. **(B)** quantification of the relative N_m_ ratio dependent on FBL. Mean ± SEM, n = 3. p = two-tailed *Student’s t-test*. **(C)** Analysis of publicly available Ribo-seq performed in HeLa cells (GSE105248) (21). This dataset contained three Ribo-seq and Input RNA-seq from FBL knockdown (Doxycycline-inducible shRNA against FBL) and control groups. Translation efficiency was calculated as normalized RPF counts divided by normalized mRNA counts. We compared genes with mapped N_m_[+1] at an upstream AUG or near-cognate codon in the 5’UTR with those lacking any N_m_ in their 5’UTR in HeLa cells. p = Wilcoxon rank-sum test. **(D)** Western blot validation of NEK2, GNL3, and SRSF1 expression upon knockdown of FBL. Representative of n=4. **(E)** quantification of the relative protein levels dependent on FBL knockdown from the gels shown (D). Mean ± SEM, n = 4. p = two-tailed *Student’s t-test*. **(F)** distribution of 2’-O-methylation stoichiometry in position N_m_[-1] of near-cognate codons within human 5’UTRs.

## DATA AVAILABILITY

This study did not generate original sequencing or structural data. Publicly available data used in this study include GEO: GSE162043 (8), GEO: GSE208837 (17), GEO: GSE105248 (21), PDB: 7UCJ (8), PDB: 6YAL (9), PDB: 8PJ1 (22), PDB: 8PJ2 (22), PDB: 8PJ3 (22), PDB: 8PJ4 (22), PDB: 8PJ5 (22), PDB: 8PJ6 (22).

